# Life histories decide reserve benefits in transient yields and bycatch persistence

**DOI:** 10.1101/2020.11.14.382705

**Authors:** Renfei Chen, Chengyi Tu, Quan-Xing Liu

**Author notes:** Corresponding author: Renfei Chen.

## Abstract

Recent research indicates that marine reserves can both improve fisheries yields of target species and maintain the persistence of bycatch species. However, the prevalent equilibrium analyses prevent our understandings in transient behavior at short-time scales. Here, we develop high dimensional theoretical frameworks by considering age structure to assess the relative advantages between reserve-only and no-reserve fisheries management strategies. Our results show that whether strategies with only reserves can achieve higher fisheries yields (measured by both weight and number) and maintain bycatch persistence depends on the life histories of both target and bycatch species through perspectives of transient oscillations. Our research has important practical applications especially for the West Coast groundfish fishery in the USA, as it suggests that reserves can perform benefits in both fisheries and conservation goals for target species with older ages at maturity and lower adult survivorship.

## Introduction

A central problem in marine ecology is improving harvested yields in fisheries and maintaining species persistence and biodiversity in conservation. To meet these goals simultaneously, different fisheries strategic managements have been proposed (Game *et al.* 2009; Gaines *et al.* 2010; Cohen & Foale 2013; Chen *et al.* 2020). Among which the most influenced methods are traditional fixed/limitation harvesting strategies and relatively new proposed ones by designing marine reserves (Hastings & Botsford 1999; Hastings *et al.* 2017; Hilborn 2017). The inevitable problem deserving attention is that which method can perform better to achieve higher fisheries yields, but without facing the risk of species extinction because of overfishing. A theoretical framework suggests that establishing marine reserves is a better choice by achieving equivalent fisheries yields with advantages in maintaining species persistence in comparison with traditional fisheries management by harvesting effort control each year (Hastings & Botsford 1999). However, the single species assumption prevents more general predictions in biodiversity such as considering the effect of some more easily extinct species relative to the target species in fisheries.

The incidental catch of unwanted species (i.e. the so-called “bycatch species”) is ubiquitous and has been a great threat to fisheries sustainability (Aalto & Baskett 2013; Komoroske & Lewison 2015; Scales *et al.* 2018; Welch *et al.* 2018). Therefore, exploring strategies to improve the harvested yields of the target species but keeping the persistence of the bycatch species have great meanings in the practical applications. To date, protecting the bycatch species and simultaneously optimizing the target fisheries yields are still a challenge due to different sustainable catch rates among species (Hastings *et al.* 2017). Nevertheless, an extended two-species model predicts that establishing marine reserve can achieve conservation and fisheries benefits simultaneously, especially when the bycatch species are long-lived with low fecundity (Hastings *et al.* 2017). Although existing multispecies with mixed life histories demonstrates the important role that marine reserve played in ecosystem-based fisheries management, more specific information about how life histories affect the relative advantages between different fisheries management methods (e.g. traditional fishing effort control vs. establishing marine reserves) still needs further answers. Besides, previous researches suggest that a model with age structure plays an important and non-negligible role in deciding which fisheries management method is better (Hilborn 2017). Moreover, the limitations of implicit equilibrium conditions prevent barriers to investigate how the advantages of marine reserves in improving fisheries yields and maintaining bycatch persistence vary in relatively short time scales before the ecosystem achieves stable equilibrium states; that is the insight into the ecosystem with transient dynamics (Hopf *et al.* 2016; Hastings *et al.* 2018; Morozov *et al.* 2019).

Understanding how transient dynamics of fisheries yields response to different management strategies is essential for monitoring fisheries and making the right decisions in adaptive management. Adaptive management is a widely accepted iterative approach to frequently regulate fisheries policy implementation based on the difference between empirical monitoring data and theoretical expectations (Kaplan *et al.* 2019; Nickols *et al.* 2019). It has been suggested that setting an expected timeline is necessary for adaptive management during the recovery of harvested populations (Kaplan *et al.* 2019). This derives from the fact that, once marine reserves are established, the abundance of fish populations may vary greatly and lead to undetectability in fisheries caused by a variety of factors including life histories (White *et al.* 2013; Kaplan *et al.* 2019; Nickols *et al.* 2019). These findings provide important inspirations to compare the relative yields advantages between fisheries management strategies of traditional effort control and the implementation of marine reserves. If the transient oscillations are too strong and the amplitude is big enough, fisheries yields under reserve implementation policy can probably be higher at one moment but lower at the other moment relative to the yields using nonspatial approaches, which increases the difficulty to decide between different fisheries management strategies. It has been demonstrated that life histories can greatly regulate transients in population abundance, and fish individuals with old ages at maturity and low natural mortality rates will increase transient dynamics (White *et al.* 2013).

To explore the transient phenomena in fisheries management as well as the intrinsic mechanism, we develop theoretical frameworks consist of reserve-only and no-reserve models which are derived from the two extreme cases in previous models (Hastings *et al.* 2017). We further extend the models to high dimensions by adding age structure to investigate the transient behavior of the target fish species (who are easily persistent even under the stress of fishing) under the persistence of the bycatch species (who have a high risk of going extinct) based on several assumptions. First, we assume that the transient dynamics of the target fish species are independent of the dynamics of the bycatch species (i.e. no ecological connections between two species) (Chen 2020). Second, we assume that the movement abilities are different between fish adults and larvae. Adults tend to be sedentary while larvae can disperse everywhere. Third, we assume that the proportions of fish individuals that are subject to reserve protection (i.e. marine reserve size) in the reserve-only model are the same for both target and bycatch species, and the fractions of fish stocks that left unharvested (i.e. escapement rate) in the no-reserve model are the same for two species. This assumption is possible in the real world when the costs of making fishing gear selective are relatively high. With these assumptions, we compare the fisheries yields measured by both number and weight under two different fisheries management strategies and explore the mechanism that causes the variations of the relative fisheries yields advantages. We investigate the effect of life histories on transient metrics through sensitive analysis, and wavelet analysis (which is an accepted method in transient analyses in ecology; see (Torrence & Compo 1998; Cazelles *et al.* 2008; Cazelles *et al.* 2014)) is used to observe the periodicity of the transient dynamics of the target fished species. With these analyses, we aim to answer three questions: i) In transient time scales, whether and when the reserve-only policy can achieve advantages of fisheries yields under bycatch species persistence relative to no-reserve policy in fisheries management? ii) Whether transient fisheries management strategies depend on different fisheries yields measurements (i.e. number vs. weight)? iii) How life histories of both target fish species and bycatch species regulate the relative fisheries yields advantages between two fisheries management strategies during transient dynamics?

## Model

### Model overview

We study two-species systems with both the reserve-only model and the no-reserve model, which can achieve equivalent fisheries yields based on previous conclusions (Hastings & Botsford 1999; Hastings *et al.* 2017). The two species are the target species with strong stock in fisheries whose life histories such as high fecundity, early maturity make is easy to persist and the bycatch weak stock species which has reverse life histories and easier to reach an unacceptably low level. We consider the age structure (results without age structure are shown in Fig. S1) of the target species while ignoring the age structure of the weak stock species so that to simplify and focus on the central issue in this study. In addition to increasing age for all adults each year, the age structure of the target species is featured with two points: i) all the youngest adults come from the larvae recruitment in one year; ii) all the oldest adults (i.e. reach the maximum age) will die next year. Among different age classes, we assume that older mature individuals have higher fecundity. See schematic in Fig. 1.

**Figure 1.**
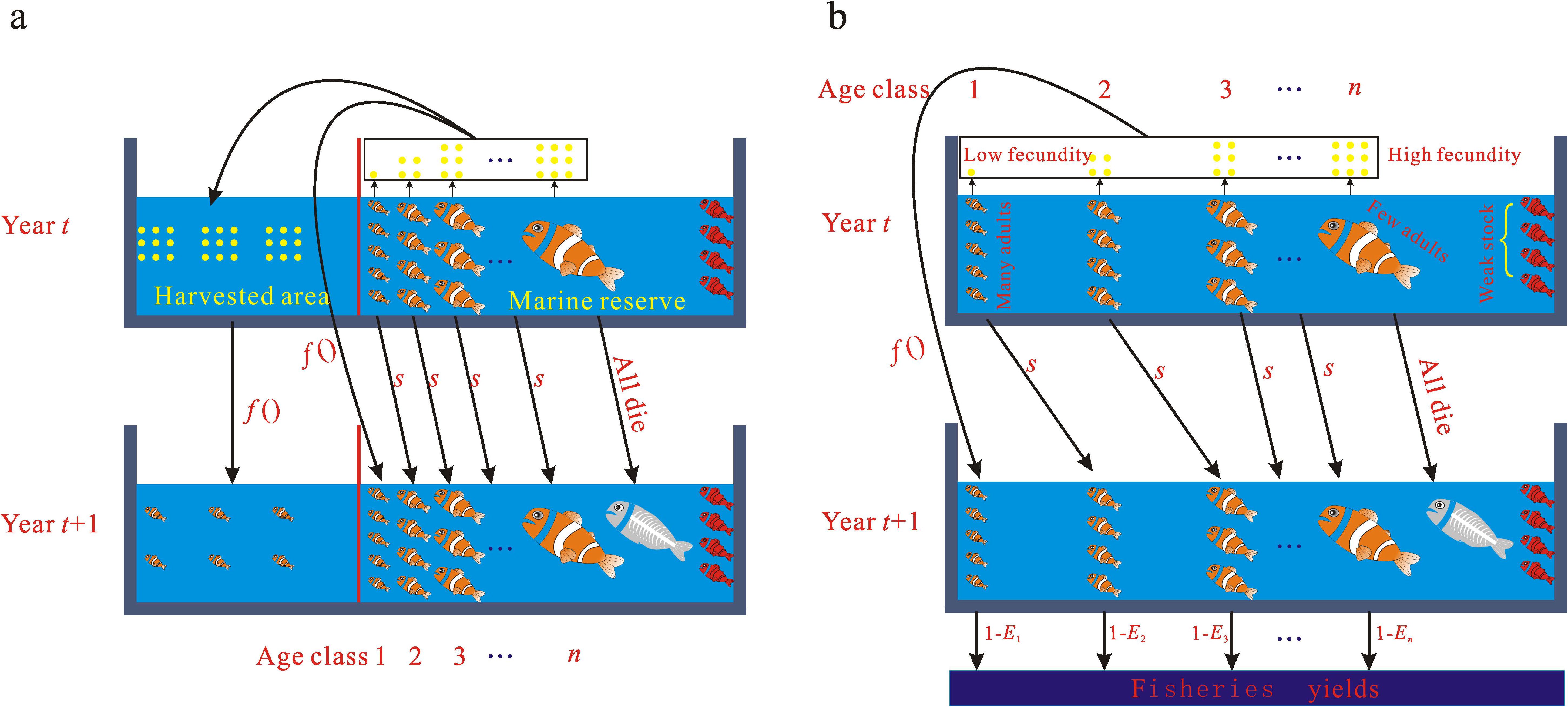
Schematic of the theoretical frameworks. (a) fisheries strategies by only using reserves. (b) traditional fisheries strategies by fishing effort control without reserves. The life histories of both target and bycatch species do not change between different fisheries strategies.

The per-capita fecundity in each age class is expressed as ***m** = (m_l_, m_2_,…, m_n_)^T^*, and the corresponding weight per individual is ***B** = (B_1_, B_2_, …, B_n_)^T^*. We denote *a_M_* as the age at maturity, then fecundity is zero when fish individual age is smaller than the threshold *a_M_* and the fecundity (expressed as a function of length) 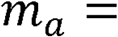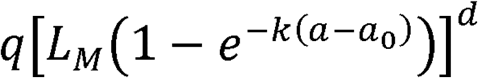 is when fish individual age surpasses the threshold of maturity (White *et al.* 2013; Kaplan *et al.* 2019), where *a* is fish individual age, *L_M_* is the fish individual asymptotic maximum length, *k* is the growth rate, *q* and *d* are constants, and *a*_0_ is the age at length zero. Similarly, fish individual weight at age *a* is expressed as 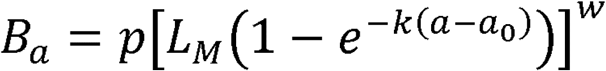, where *p* and *w* are constants (Kaplan *et al.* 2019).

### Part 1 reserve-only model

When only using marine reserves rather than fixed harvesting rates as fisheries management methods, an implicit assumption is that all adult fish are harvested outside marine reserves. To maintain the persistence of weak stock species (i.e. non-target fish species), the reserve size *c* should be no smaller than a critical value. According to previous research (Hastings *et al.* 2017), the critical value is

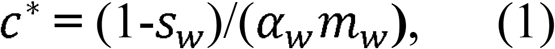

where *s_w_*, *α_w_*, and *m_w_* are survival rate, proliferation rate per generation, and per-capita fecundity for the weak stock species, respectively.

The adult density with age structure at time *t* inside the marine reserve is defined as 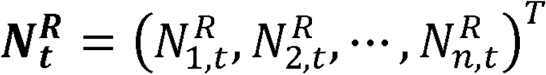. Thus, the total number of larvae produced inside the marine reserve is the dot product of adult density and fecundity multiplied by the reserve size. Based on the “well-mixed” assumption as suggested in previous research (Hastings & Botsford 1999; Hart 2006; Hopf *et al.* 2019), the larvae density both inside and outside the marine reserve is 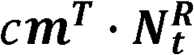 which equals the total number because the total marine system size is scaled as 1. Therefore, the density of larvae that survive to adults will be

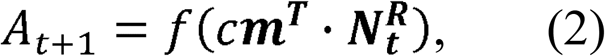

where *f*(·) is the Beverton–Holt growth function. We further define the adult natural survival rate each year as a fixed value *s*; note that all fish individuals will die next year once they achieve the maximum age (See Fig. 1a). According to simple derivations (see Section one in appendix), the age-structured adult density inside reserve at time *t* is

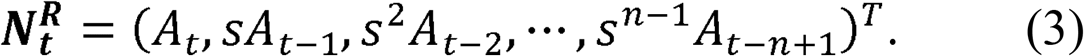

The fisheries yields measured by number can be expressed as

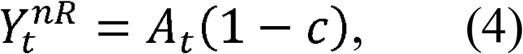

and the fisheries yields measured by weight can be expressed as

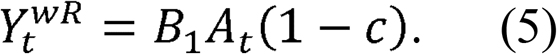

### Part 2 no-reserve model

When using traditional ways of fixed harvesting rate without marine reserves (See Fig. 1b), the escapement rate *E* should be no smaller than the critical value *E*^*^ to maintain the persistence of the weak stock species. Based on previous research (Hastings *et al.* 2017), we have

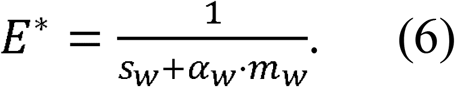

Defining *a_E_* as the critical age that fish individuals are harvested, the escapement rate is 1 when fish individual age is smaller than *a_E_*, and the escapement rate is *E*^*^ when fish individual age is larger than *a_E_*. With a general form, we denote the escapement rate for the age-structured strong stock species as ***E** = (E_1_, E_2_, …, E_n_)^T^*, and the population density at time *t* is 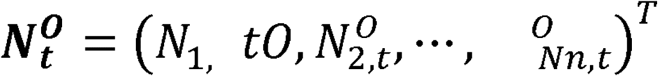. Similar to the reserve-only model, the total larvae produced by all the adults are the dot product of adult density and fecundity. Thus, the density of larvae that survive to adults will be

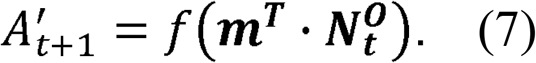

The age-structured adult density at time *t* is

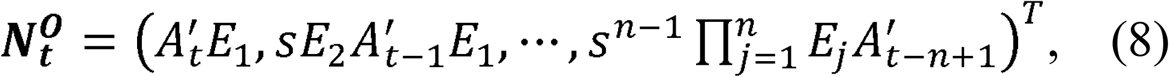

and the specific derivation can be seen in Section one in the appendix. To exhibit fisheries yields with concise form in the next step, we show the density at time *t* before harvesting as

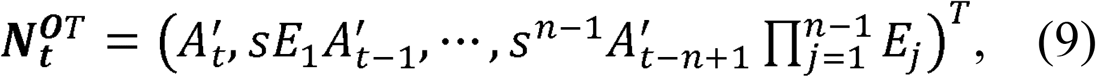

and thus, the fisheries yields in number can be expressed as

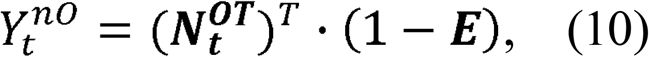

and the fisheries yields in weight can be expressed as

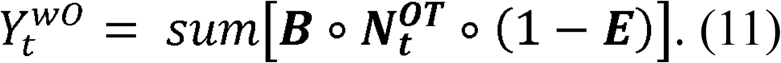

## Results

Our results have shown the transient oscillations can occur for the dynamics of population densities (Fig. 2a_1_-a_7_), fisheries yields measured by number (Fig. 2b_1_-b_7_) and fisheries yields measured by weight (Fig. 2c_1_-c_7_), and the periodicity is exhibited with wavelet analyses (Fig. 3). Moreover, regardless of the measurement methods (i.e. measured by weight or measured by number), the fisheries yields produced by the reserve-only model can always be higher (red) or lower (yellow) than the yields produced by the no-reserve model with appropriate hypothetical parameter settings during the whole transient period (Fig. 2b_1_; Fig. 2c_1_). In other cases, it would be difficult to judge which model can achieve higher fisheries yields as the transient oscillations lead to higher fisheries yields with the reserve-only model at one moment while lower ones at another moment and this is true for both kinds of yields measured by weight and by number (blue lines in Fig. 2b_1_ and Fig. 2c_1_). The parameter values are arbitrary. However, the yield comparison conclusion that three situations (i.e. fisheries yields with the reserve-only model is higher, lower, or difficult to judge relative to yields with no-reserve model) can happen is robust with other parameters values and with different bycatch species for all the hypothetical cases (Fig. 2b_1_-b_3_; Fig. 2c_1_-c_3_). For the empirical cases, the results about the relative advantages of fisheries yields between two fisheries management methods (i.e. methods with reserve-only model vs. no-reserve model) are consistent with that for the hypothetical cases (i.e. all three situations can occur) if fisheries yields are measured by weight (Fig. 2c_4_-c_7_). However, if fisheries yields are measured by the number, fisheries yields with the reserve-only model are always higher relative to that with the no-reserve model (Fig. 2b_4_-b_7_).

**Figure 2.**
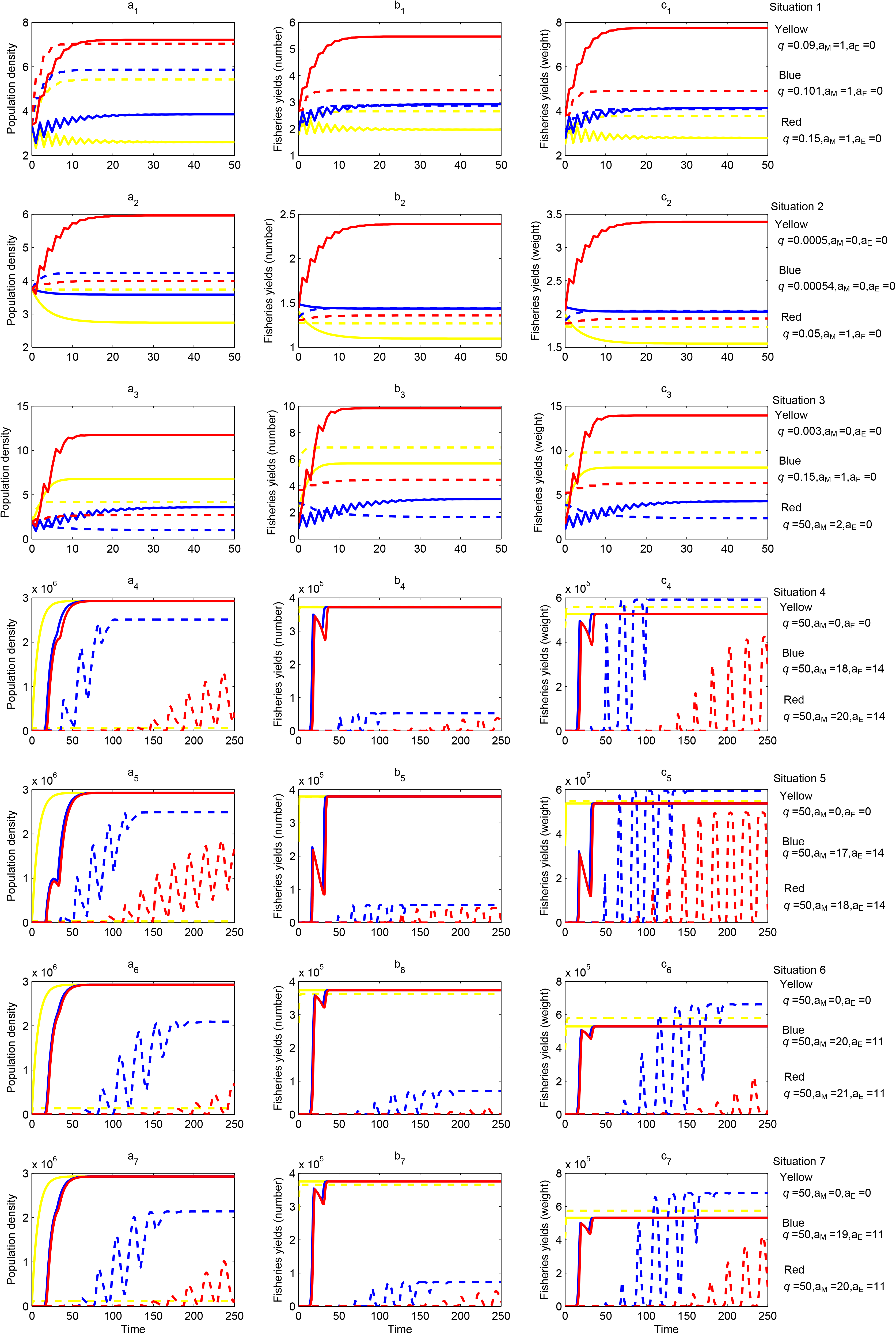
Transient dynamics of population density and fisheries yields for target species under the persistence of different bycatch species. Solid lines denote reserve-only strategies, and dashed lines denote no-reserve strategies. Different colors represent different parameter settings to perform sensitivity analyses. (a_1_-a_7_) Transient dynamics of population density. Note that population density with reserve-only strategies denotes density inside marine reserves. (b_1_-b_7_) transient dynamics of fisheries yields measured by number. (c_1_-c_7_) transient dynamics of fisheries yields measured by weight. a_1_-a_7,_ b_1_-b_7,_ and c_1_-c_7_ correspond to situations 1-7, and specific definitions and differences among different situations can be seen in (Chen 2020).

**Figure 3.**
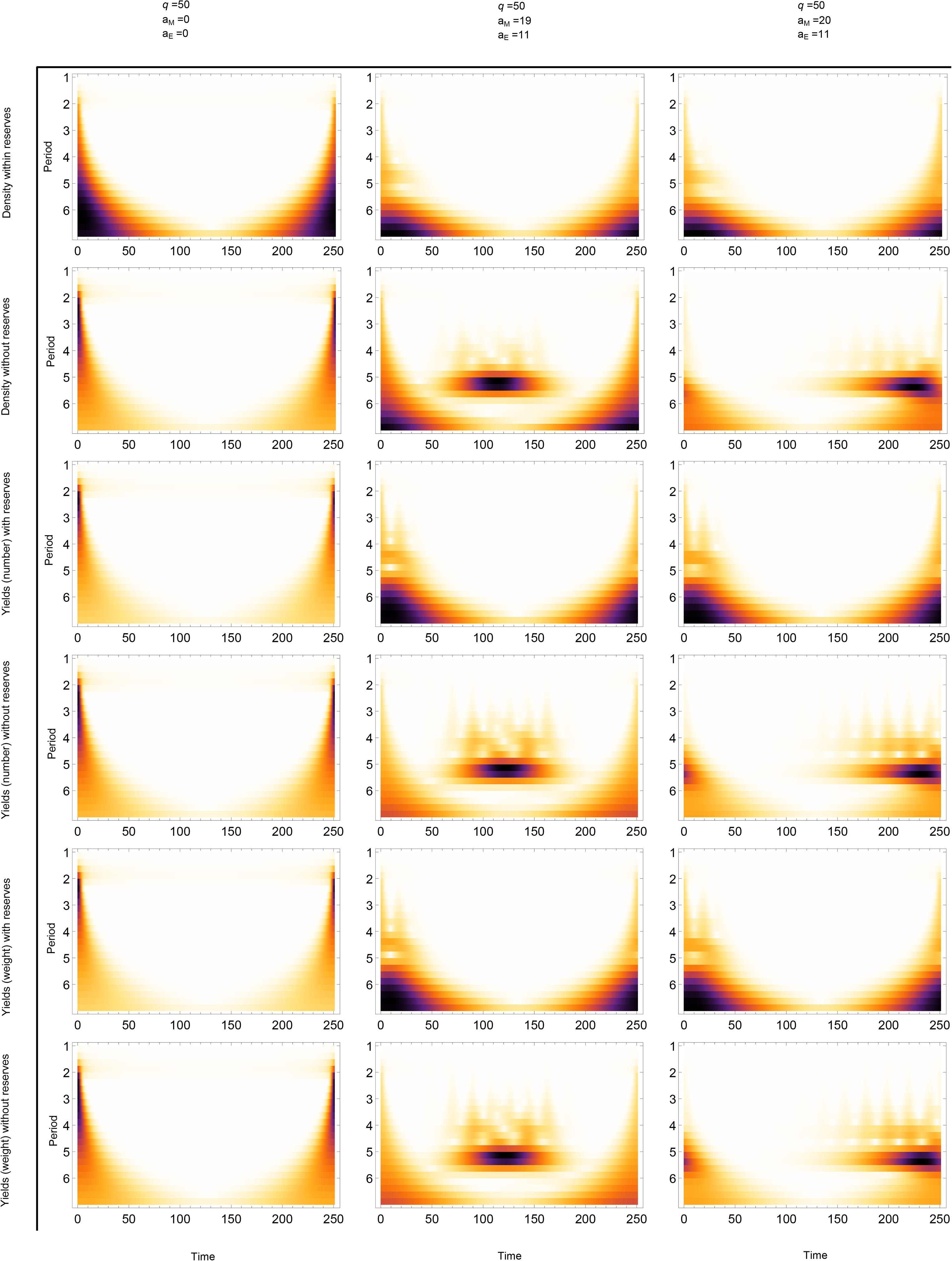
Wavelet analyses of the transient dynamics in an empirical case. (Corresponding to the last line in figure 2, i.e. situation 7). It denotes the wavelet power spectrum of the raw series, and the power values increase from white (low values) to dark red (high values).

The effect of life histories for target species on fisheries yields shows that, with empirical parameter settings, fisheries management methods with the reserve-only model can always lead to higher fisheries yields relative to the no-reserve model no matter how influencing factors (age at maturity, age at fishing and adult survivorship for the target species) varies among the suitable parameter value ranges (Fig. 4a-c). However, the relative advantages in fisheries yields change with influencing factors if fisheries yields are measured by weight (Fig. 4e-g). Specifically, the target species with life histories of younger age at maturity, younger age at fishing, and higher adult survivorship can achieve higher fisheries yields with no-reserve fisheries management method relative to the reserve-only method. On the contrary, the target species with reversed life histories can lead to higher fisheries yields with the reserve-only method relative to the no-reserve method. The robust results still hold among different weak species (shown in different colors in Fig. 4) and among different time steps (Fig. S2, S3). For the hypothetical cases, the effects of adult survivorship on fisheries yields measured by weight (i.e. higher adult survivorship benefits for methods with no-reserve model) are similar to the results for the empirical cases (Fig. 4g; Fig. S4g, S5g, S6g), while the effects of other factors (age at maturity and age at fishing) on both kinds of fisheries yields as well as the effects of adult survivorship on fisheries yields measured by number are different from that for the empirical cases (Fig. 4; Fig. S4, S5, S6).

**Figure 4.**
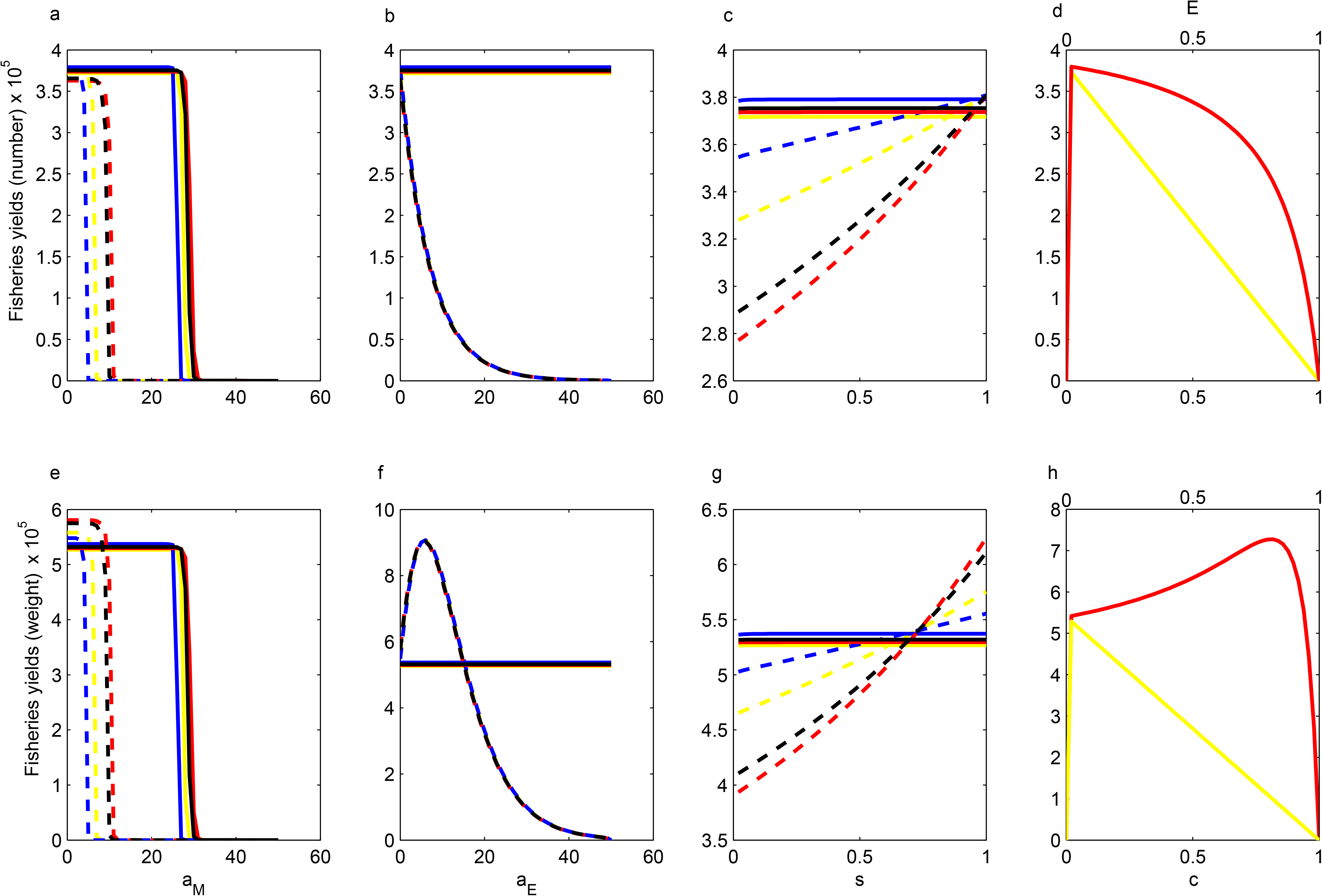
Effects of life histories and harvesting management on fisheries yields with both reserve-only and no-reserve strategies for empirical cases (situations 4-7). (a-c) Fisheries yields measured by number response to age at maturity, age at fishing, adult survivorship, respectively for target species. (d) Fisheries yields measured by number response to the critical reserve size and the critical escapement rate. (e-g) Fisheries yields measured by weight response to age at maturity, age at fishing, adult survivorship, respectively for target species. (h) Fisheries yields measured by weight response to the critical reserve size and the critical escapement rate. In a-c and e-g, solid lines denote reserve-only strategies, and dashed lines denote no-reserve strategies. Different colors represent different bycatch species. In d and h, yellow colors denote critical reserve size with reserve-only strategies, and red colors denote critical escapement rate with no-reserve strategies. All subplots denote the time step at 50. Parameter settings: (a, e) *q* = 50, a_E_ = 0; (b, f) *q* = 50, a_M_ = 1; (c, d, g, h) *q* = 50, a_M_ = 1, a_E_ = 0; see Table S1 and Materials and methods for other parameters.

For the empirical cases, fisheries yields measured by both number and weight are always higher with the no-reserve fisheries management method than that with the reserve-only method if the critical values of reserve size for the weak species persistence in the reserve-only model are identical to the critical values of escapement rate in the no-reserve model (Fig. 4d, h). However, fisheries management with the reserve-only method can achieve higher fisheries yields relative to the no-reserve method if the critical reserve size is small while the critical escapement rate is very large. On the contrary, for the hypothetical cases, fisheries yields with the no-reserve method are always lower than that with the reserve-only method when the critical reserve size and escapement rate are equal, and could be higher when the critical escapement rate is mediate and the critical reserve size is too big or too small (Fig. S4d, h; Fig. S5d, h; Fig. S6d, h). This suggests that the critical reserve size in the reserve-only model and the critical escapement rate in the no-reserve model can greatly regulate the relative yield advantages between two fisheries management methods (i.e. which method can achieve higher fisheries yields), which is important as the critical values of both reserve size and escapement rate are linked to the life histories of the bycatch weak stock species to meet the persistence condition.

Sensitive analyses show that the transient metric *θ* oscillates and has strong periodicity with the variations of influencing factors (i.e. age at maturity, age at fishing, and adult survivorship) (Fig. 5, Fig. S7, Fig. S8). It suggests that the initial trajectories of the system periodically close to or leave the stable level, and thus the amplitude of the oscillations of the population dynamics decrease or increase periodically by regulating the life histories of the target species such as age at maturity, age at fishing as well as adult survivorship.

**Figure 5.**
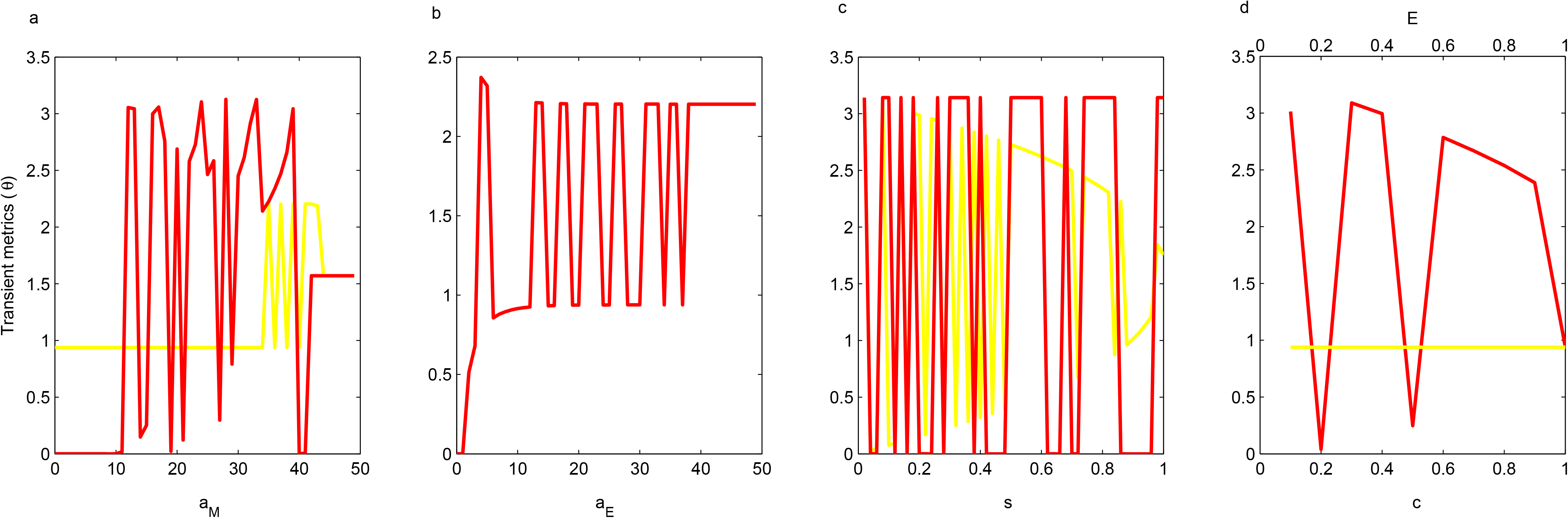
Sensitive analyses of transient metric *θ* in response to different parameters for empirical case (situation 7) with different fisheries management strategies. (a) age at maturity, (b) age at fishing, (c) adult survivorship, (d) critical reserve size (yellow) and critical escapement rate (red). In a and c, yellow color denotes reserve-only strategies, and red colors denote no-reserve strategies. Parameter settings: (a) *q* = 50, a_E_ = 0; (b) *q* = 50, a_M_ = 1; (c, d) *q* = 50, a_M_ = 1, a_E_ = 0; see Table S1 and Materials and methods for other parameters.

## Discussion

Marine reserves are important in fisheries management (Mangel 2000; Edgar *et al.* 2014; Herrera *et al.* 2016). Our results have shown that fisheries management strategies by only establishing marine reserves without fishing effort control cannot always be the optimal method to improve the fisheries yields as well as maintain species persistence if the dynamics of the two-species system is investigated from the transient short time scale rather than the asymptotic equilibrium state. When fisheries management with marine reserves can achieve higher fisheries yields relative to traditional management with fishing effort control depends on the measurement methods of fisheries yields and the life histories of both the target species and the bycatch species. If fisheries yields are measured by fish individual numbers, establishing a marine reserve could be the primary fisheries management strategies, especially for the US West Coast groundfish fishery (Hastings *et al.* 2017; Nickols *et al.* 2019; Chen 2020). If fisheries yields are measured by weight which can make the theoretical frameworks include the difference of both fecundity and biomass for fish individuals of different ages (Kaplan *et al.* 2019), traditional fishing effort control will not be the obvious disadvantage strategies in both improving fisheries yields and maintaining species persistence. The effect of fisheries yields measurement on fisheries management strategies derives from the fact that fisheries management by establishing marine reserves only harvest relatively young adults each year because all the adult fish individuals in the harvested area come from the larvae dispersal from the marine reserves, while fisheries management by fishing effort control can harvest relatively old adult which have relatively large biomass (or weight). Thus, the measurement with weight benefits for the traditional fisheries management in evaluating its advantage of improving fisheries yields.

Further mechanisms show that whether the fisheries management strategies of no-reserve can improve fisheries yields under bycatch persistence depends on the life histories of both target and bycatch species. If the target species are younger at maturity or have higher adult survivorship, strategies with no-reserve can achieve higher fisheries yields relative to the target species with older age at maturity or with lower adult survivorship (Fig. 4). Nevertheless, for the hypothetical cases, the effect of age at maturity and age at fishing on both kinds of fisheries yields are complex among different weak species relative to empirical cases; that is fisheries yields with the reserve-only model are higher relative to yields with the no-reserve model for one weak species while it is lower for another weak species (Fig. 4; Fig. S4, S5, S6). The differences between the hypothetical and the empirical cases derive from the fact that adult survivorship in the hypothetical cases is very low while it is very high in the empirical cases (Hastings *et al.* 2017; Chen 2020). Thus, both the age at maturity and at fishing cannot be too old in hypothetical cases. Otherwise, the target strong stock species cannot persist.

The life histories of the bycatch species can regulate the relative yield advantages between two fisheries management strategies through deciding the critical reserve size and escapement rate to maintain persistence according to the weak stock persistence conditions shown in Eq. 1 and Eq. 6. For example, if the critical reserve size is 0.1 and the critical escapement rate is 0.95 so that fisheries management with reserve-only strategies can achieve higher fisheries yields relative to no-reserve strategies (Fig. 4), the corresponding life histories of the bycatch species can be predicted by the combination of Eq. 1 and Eq. 6 through controlling one life-history parameter (e.g. achieve adult survivorship and fecundity by controlling proliferation rate per generation). On the contrary, solving equations 1 and 6 with the assumption that the critical reserve size equals the critical escapement rate can achieve life histories of bycatch under which strategies with no-reserve produce higher fisheries yields relative to strategies with only reserves. In such a case, even if the bycatch species are long-lived with low fecundity which is important based on the previous model (Hastings *et al.* 2017), reserves cannot make sure benefits simultaneously in fisheries and conservation.

The life histories can regulate the relative fisheries yields advantages between two fisheries management strategies through the other perspective by regulating the amplitude of the transient oscillations. To make the relative yield advantages predictable, the amplitude of the transient oscillations of the system should not be too big. Otherwise, the “up and down” of the fisheries yields with time will make it complex to predict which fisheries management strategy (no-reserve vs. reserve-only) is more suitable for both conservation and fisheries goals. To achieve small amplitude of the transient oscillations, the life histories of both the target and bycatch species can be regulated because of the periodical relationships between the transient metric *θ* (which decides the amplitude of transient oscillations (White *et al.* 2013; Kaplan *et al.* 2019; Chen 2020)) and life history parameters of both the target and bycatch species (Fig.5).

We consider the effect of age structure in the main research. If there is no age structure as shown in Hastings et al. (2017) (fisheries yields measured by weight are identical to that measured by number), the transient analysis suggests that traditional fisheries management methods can achieve higher fisheries yields relative to strategies with marine reserves, but it cannot maintain the persistence of the bycatch species (yellow color in Fig. S1). When bycatch species is persistent, strategies with marine reserves can achieve higher fisheries yields (Fig. S1), which is consistent with previous equilibrium research that strategies with marine reserves can perform an important role in fisheries management by solving bycatch problems (Hastings *et al.* 2017). However, when the adult survivorship is very high as shown in the empirical cases, traditional strategies without reserves can achieve higher fisheries yields relative to strategies with marine reserves and maintain bycatch species persistence at the beginning of the transient time scales, which is consistent with our results with age structure regardless of the results without age structure at the equilibrium state still show the advantage of strategies with marine reserves (Fig. 2; Fig. S1h, j, l, n). Besides, relative to our theoretical frameworks, models developed by Hastings et al. (2017) do not show obvious transient phenomena (Fig. S1), which are consistent with previous conclusions that transients are difficult to observe in models without age structure (White *et al.* 2013; Chen 2020). This derives from the fact that age structure often leads to high dimensions which have been suggested to be the main mechanism to cause transients (Hastings *et al.* 2018; Morozov *et al.* 2019).

Previous investigations in the transient dynamics of fish abundance after establishing marine reserves indicate that the transient metric *θ* (which determines the amplitude of the transient oscillations) becomes, in general, large with the increasing of age at maturity and small with the increasing of natural mortality (White *et al.* 2013). However, our sensitivity analyses indicate that the transient metric *θ* increases and decreases periodically with the variation of both ages at maturity and adult survivorship (Fig. 5). The difference between our research and previous ones (White *et al.* 2013) derives from the fact that previous research considers the transient response of linear models with Leslie matrices while we consider the non-linear ones and thus the eigenvalues of matrix ***A*** and ***A***′ vary with population density (Section two in appendix). These results suggest that considering the non-linear effect of larval survival to adult will increase the difficulty of investigating the transient dynamics as the predictability periodically varies with life histories, and thus increase the difficulty to judge which fisheries management strategies (reserve-only vs. no-reserve) perform better. Further insight into the periodicity of the transient dynamics of the system with time is helpful for predictability, and our wavelet analyses provide answers for this (Fig. 3; Fig. S9-S14).

Our researches are based on the “scorched earth” assumption (Malvadkar & Hastings 2008; Edwards *et al.* 2010) for the fisheries management with reserve-only strategies, which does not consider the effect of economic costs in fisheries(Sanchirico *et al.* 2006; White *et al.* 2008). If the effect of economic costs is considered, harvesting all adults in the harvested area (i.e. the scorched earth assumption) can lead to high harvesting costs in fisheries as the catchability decreases with the decreasing of population density in the natural system. In that case, costs with reserve-only strategies are higher and may not be a better choice for fisheries management relative to traditional methods without reserves.

We ignore the age structure of the bycatch species so that the persistence conditions are easy to achieve although the conditions are necessary but not sufficient (Chen 2020). By considering the age structure of the bycatch species, high dimensions lead to much complex persistence conditions, which deserves developing new analytical and simulation approaches. However, similar conclusions may be achieved even if we consider the effect of the age structure of the bycatch species if new theoretical frameworks are still based on the assumption that there are no ecological connections between the target and the bycatch species. Future works can explore the predator-prey or competition connections between the target and the bycatch species, which can greatly regulate the population density of both species (Frid & Marliave 2010; Aalto & Baskett 2013) and thus further play a role in predicting relatively higher fisheries yields between different fisheries management strategies under bycatch persistence in transient time scales (specific information is shown in another manuscript).

## Materials and methods

With both reserve-only and no-reserve models, we first simulate the transient dynamics of the population density and fisheries yields of the target species. The system starts with the initial condition at a stable age distribution. To make a comparison, the initial conditions are the same for both reserve-only and no-reserve models. The parameter values for simulations come from previous researches (Hagerman 1952; Hunter *et al.* 1990; White *et al.* 2013; Hastings *et al.* 2017; Kaplan *et al.* 2019; Chen 2020) and can be seen in Table S1. We use the average values if these parameters are different between male and female individuals. Specifically, *a*_0_ = −5.10, *d* = 2.96, *p* = 2.41 x 10^−4^, *w* = 2.96, *k* = 0.087, *L_M_* = 45.55, and the maximum age is 50. Coefficient *q* is regulated to maintain a positive growth rate. The Beverton–Holt functional form is 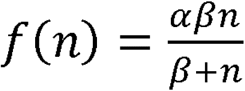 (Chen 2020), where *α* and *β* denote the proliferation rate per generation and carrying capacity, respectively. We explore the effect of life histories on transient metric *θ* which determines the amplitude of transient oscillations and measures how close the initial system state is to the final equilibrium state (White *et al.* 2013; Kaplan *et al.* 2019). To calculate transient metric *θ*, we transform both reserve-only and no-reserve models into identical cases (see details in Section two in appendix).

## Supporting information

Model derivation and figures in appendix

## Ethics

This article has no ethical problems.

## Data accessibility

This article has no additional data and there is no need to deposit data to a public repository.

## Author contributions

Renfei Chen designed research, performed research, and wrote the paper; Chengyi Tu performed the wavelet analyses; Quan-Xing Liu edited and wrote the paper.

