## Supplementary material for "Life histories decide reserve benefits in transient yields and bycatch persistence": Model derivation and figures in appendix

### Section one. Derivation of population density with reserve-only and no-reserve models

Based on our definitions in the main text, the adult density at time *t* inside the marine reserve is $\boldsymbol{N_t^R} = \left(N_{1,t}^R, N_{2,t}^R, \cdots, N_{n,t}^R\right)^T$ in the reserve-only model, and the density of larvae that survive to adults is $A_{t+1} = f(c\boldsymbol{m^T} \cdot \boldsymbol{N_t^R})$. Thus, the density inside reserves of each age class from year 1 to year *t* can be expressed as

$$\boldsymbol{N_1^R} = \left(N_{1,1}^R, N_{2,1}^R, \cdots, N_{n,1}^R\right)^T$$

$$\boldsymbol{N_2^R} = \left(A_2, sN_{1,1}^R, sN_{2,1}^R, \cdots, sN_{n-1,1}^R\right)^T$$

$$\boldsymbol{N_3^R} = \left(A_3, sA_2, s^2N_{1,1}^R, s^2N_{2,1}^R, \cdots, s^2N_{n-2,1}^R\right)^T$$

$$\boldsymbol{N_4^R} = \left(A_4, sA_3, s^2A_2, s^3N_{1,1}^R, s^3N_{2,1}^R, \cdots, s^3N_{n-3,1}^R\right)^T$$

$$\vdots$$

$$\boldsymbol{N_n^R} = \left(A_n, sA_{n-2}, \cdots, s^{n-2}A_2, s^{n-1}N_{1,1}^R\right)^T$$

$$\boldsymbol{N_{n+1}^R} = (A_{n+1}, sA_n, s^2A_{n-1}, \cdots, s^{n-1}A_2)^T$$

$$\boldsymbol{N_{n+2}^R} = (A_{n+2}, sA_{n+1}, s^2A_n, \cdots, s^{n-1}A_3)^T$$

$$\boldsymbol{N_{n+3}^R} = (A_{n+3}, sA_{n+2}, s^2A_{n+1}, \cdots, s^{n-1}A_4)^T$$

$$\vdots$$

$$\boldsymbol{N_{n+i}^R} = (A_{n+i}, sA_{n+i-1}, s^2A_{n+i-2}, \cdots, s^{n-1}A_{i+1})^T$$

Let $n+i=t$, then $i=t-n$. So,

$$\boldsymbol{N_t^R} = (A_t, sA_{t-1}, s^2A_{t-2}, \cdots, s^{n-1}A_{t-n+1})^T$$

Note that the transient time *t* is larger than the age class *n*. Similarly, in no-reserve model, the adult density of each age class from year 1 to year *t* can be expressed as

$$\boldsymbol{N_1^O} = \left(N_{1,1}^O, N_{2,1}^O, \cdots, N_{n,1}^O\right)^T$$

$$\boldsymbol{N_2^O} = \left(A_2' E_1, sE_2 N_{1,1}^O, sE_3 N_{2,1}^O, \cdots, sE_n N_{n-1,1}^O\right)^T$$

$$\boldsymbol{N_3^O} = \left(A_3' E_1, sE_2 A_2' E_1, sE_3 sE_2 N_{1,1}^O, \cdots, sE_n sE_{n-1} N_{n-2,1}^O\right)^T$$

$$\boldsymbol{N_4^O} = \left(A_4' E_1, sE_2 A_3' E_1, sE_3 sE_2 A_2' E_1, \cdots, sE_n sE_{n-1} sE_{n-2} N_{n-3,1}^O\right)^T$$

$$\vdots$$

$$\boldsymbol{N_n^O} = \left(A_n' E_1, sE_2 A_{n-1}' E_1, \cdots, s^{n-2} \prod_{j=1}^{n-1} E_j A_2', s^{n-1} \prod_{j=2}^{n} E_j\, N_{1,1}^O\right)^T$$

$$\boldsymbol{N_{n+1}^O} = \left(A_{n+1}' E_1, sE_2 A_n' E_1, \cdots, s^{n-1} \prod_{j=1}^{n} E_j A_2'\right)^T$$

$$\boldsymbol{N_{n+2}^O} = \left(A_{n+2}' E_1, sE_2 A_{n+1}' E_1, \cdots, s^{n-1} \prod_{j=1}^{n} E_j A_3'\right)^T$$

$$\boldsymbol{N_{n+3}^O} = \left(A_{n+3}' E_1, sE_2 A_{n+2}' E_1, \cdots, s^{n-1} \prod_{j=1}^{n} E_j A_4'\right)^T$$

$$\vdots$$

$$\boldsymbol{N_{n+i}^O} = \left(A_{n+i}' E_1, sE_2 A_{n+i-1}' E_1, \cdots, s^{n-1} \prod_{j=1}^{n} E_j A_{i+1}'\right)^T$$

Let $n+i=t$, then $i=t-n$. So,

$$\boldsymbol{N_t^O} = \left(A_t' E_1, sE_2 A_{t-1}' E_1, \cdots, s^{n-1} \prod_{j=1}^{n} E_j A_{t-n+1}'\right)^T$$

**Section two. Models for transient metrics**

To perform sensitive analysis through transient metrics, our theoretical models with recurrence formula in the main text can be transformed into equivalence equations.

For the reserve-only model, we have

$$\boldsymbol{N_{t+1}^{R}} = \boldsymbol{A}\,\boldsymbol{N_{t}^{R}}\ ,$$

where the matrix **A** is expressed as

$$\boldsymbol{A} = \begin{vmatrix} A_t/N_{1,t}^R & 0 & \dots & 0 & 0 \\ s & & & & \\ & s & & & \\ & & \ddots & & \\ & & & s & 0 \end{vmatrix}$$

Therefore, we could achieve that the eigenvalues of **A** is not constant but vary with time because the elements in matrix **A** contain population density which change with transient time. The fisheries yields measured by number ($Y_t^{nR}$) and by weigh ($Y_t^{wR}$) with reserve-only model are

$$Y_t^{nR} = A_t(1-c), \text{ and}$$

$$Y_t^{wR} = B_1 A_t (1-c).$$

For the no-reserve model, we have

$$\boldsymbol{N_{t+1}^{O}} = \boldsymbol{A'}\boldsymbol{N_{t}^{O}}$$

where the matrix $\boldsymbol{A'}$ is expressed as

$$\boldsymbol{A'} = \begin{vmatrix} A_t' \cdot E_1/N_{1,t}^O & 0 & \dots & 0 & 0 \\ sE_2 & & & & \\ & sE_3 & & & \\ & & \ddots & & \\ & & & sE_n & 0 \end{vmatrix}$$

Similarly, the eigenvalues of $\boldsymbol{A'}$ vary with time. The fisheries yields measured by number ($Y_t^{nO}$) and by weigh ($Y_t^{wO}$) with no-reserve model are

$$Y_t^{nO} = (\boldsymbol{A''}\boldsymbol{N_t^O})^T \cdot (1-\boldsymbol{E}), \text{ and}$$

$$Y_t^{wO} = sum\left[\boldsymbol{B} \circ \left(\boldsymbol{A''}\boldsymbol{N_t^O}\right) \circ (1-\boldsymbol{E})\right].$$

where $\circ$ denotes Hadamard product among different vectors, and the matrix $\boldsymbol{A''}$ is expressed as

$$A'' = \begin{vmatrix} A'_t/N^O_{1,t} & 0 & \cdots & 0 & 0 \\ s & & & & \\ & s & & & \\ & & \ddots & & \\ & & & s & 0 \end{vmatrix}$$

**Table S1** Specific parameter values for analyses. See definitions and differences of situations 1-7 in (Chen 2020).

| Parameters | Situation 1 | Situation 2 | Situation 3 | Situation 4 | Situation 5 | Situation 6 | Situation 7 |
|---|---|---|---|---|---|---|---|
| s | 0.01 | 0.01 | 0.01 | 0.87 | 0.87 | 0.87 | 0.87 |
| m | 8.5 | 8.5 | 8.5 | 1 | 1 | 1 | 1 |
| $\alpha$ | 0.7 | 0.7 | 0.7 | 16 | 16 | 16 | 16 |
| $\beta$ | 20 | 20 | 20 | 23799 | 23799 | 23799 | 23799 |
| $s_w$ | 0.85 | 0.5 | 0.7 | 0.85 | 0.939 | 0.95 | 0.955 |
| $m_w$ | 1.6 | 1.4 | 2.5 | 1 | 1 | 1 | 1 |
| $\alpha_w$ | 0.4 | 0.6 | 0.78 | 6.26 | 13.62 | 2.67 | 3.14 |
| $\beta_w$ | 5 | 5 | 5 | 825.8 | 204.05 | 3495.3 | 72.59 |

**Figure legends in appendix.**

Figure S1. Transient dynamics without age structure.

Figure S2. Effects of life histories and harvesting management on fisheries yields with both reserve-only and no-reserve strategies for empirical cases (situations 4-7). All parameters are the same as that in Fig. 4 with only difference that the system runs to time step 100.

Figure S3. Effects of life histories and harvesting management on fisheries yields with both reserve-only and no-reserve strategies for empirical cases (situations 4-7). All parameters are the same as that in Fig. 4 with only difference that the system runs to time step 150.

Figure S4. Effects of life histories and harvesting management on fisheries yields with both reserve-only and no-reserve strategies for hypothetical cases (situations 1-3). The system runs to time step 10. Parameter settings are: (a, e) $q$ =0.09, $a_E$ = 0; (b, f) $q$ = 0.09, $a_M$ = 1; (c, d, g, h) $q$ = 0.09, $a_M$ = 1, $a_E$ = 0; see Table S1 and Materials and methods for other parameters.

Figure S5. Effects of life histories and harvesting management on fisheries yields with both reserve-only and no-reserve strategies for hypothetical cases (situations 1-3). All parameters are the same as that in Figure. S4 with only difference that the system runs to time step 20.

Figure S6. Effects of life histories and harvesting management on fisheries yields with both reserve-only and no-reserve strategies for hypothetical cases (situations 1-3). All parameters are the same as that in Figure. S4 with only difference that the system runs to time step 40.

Figure S7. Sensitive analyses of transient metric $\theta$ in response to different parameters for empirical cases (situations 4-6) with different fisheries management strategies. (a, e, i) age at maturity, (b, f, j) age at fishing, (c, g, k) adult survivorship, (d, h, l) critical reserve size (yellow) and critical escapement rate (red). Parameter settings: (a, e, i) $q$ = 50, $a_E$ = 0; (b, f, j) $q$ = 50, $a_M$ = 1; (c, d, g, h, k, l) $q$ = 50, $a_M$ = 1, $a_E$ = 0; see Table S1 and Materials and methods for other parameters.

Figure S8. Sensitive analyses of transient metric $\theta$ in response to different parameters for hypothetical cases (situations 1-3) with different fisheries management strategies. (a, e, i) age at maturity, (b, f, j) age at fishing, (c, g, k) adult survivorship, (d, h, l) critical reserve size (yellow) and critical escapement rate (red). Parameter settings: (a, e, i) $q$ =0.09, $a_E$ = 0; (b, f, j) $q$ = 0.09, $a_M$ = 1; (c, d, g, h, k, l) $q$ = 0.09, $a_M$ = 1, $a_E$ = 0; see Table S1 and Materials and methods for other parameters.

Figure S9. Wavelet analyses of the transient dynamics in hypothetical case (situation 1).

Figure S10. Wavelet analyses of the transient dynamics in hypothetical case (situation 2).

Figure S11. Wavelet analyses of the transient dynamics in hypothetical case (situation 3).

Figure S12. Wavelet analyses of the transient dynamics in empirical case (situation 4).

Figure S13. Wavelet analyses of the transient dynamics in empirical case (situation 5).

Figure S14. Wavelet analyses of the transient dynamics in empirical case (situation 6).

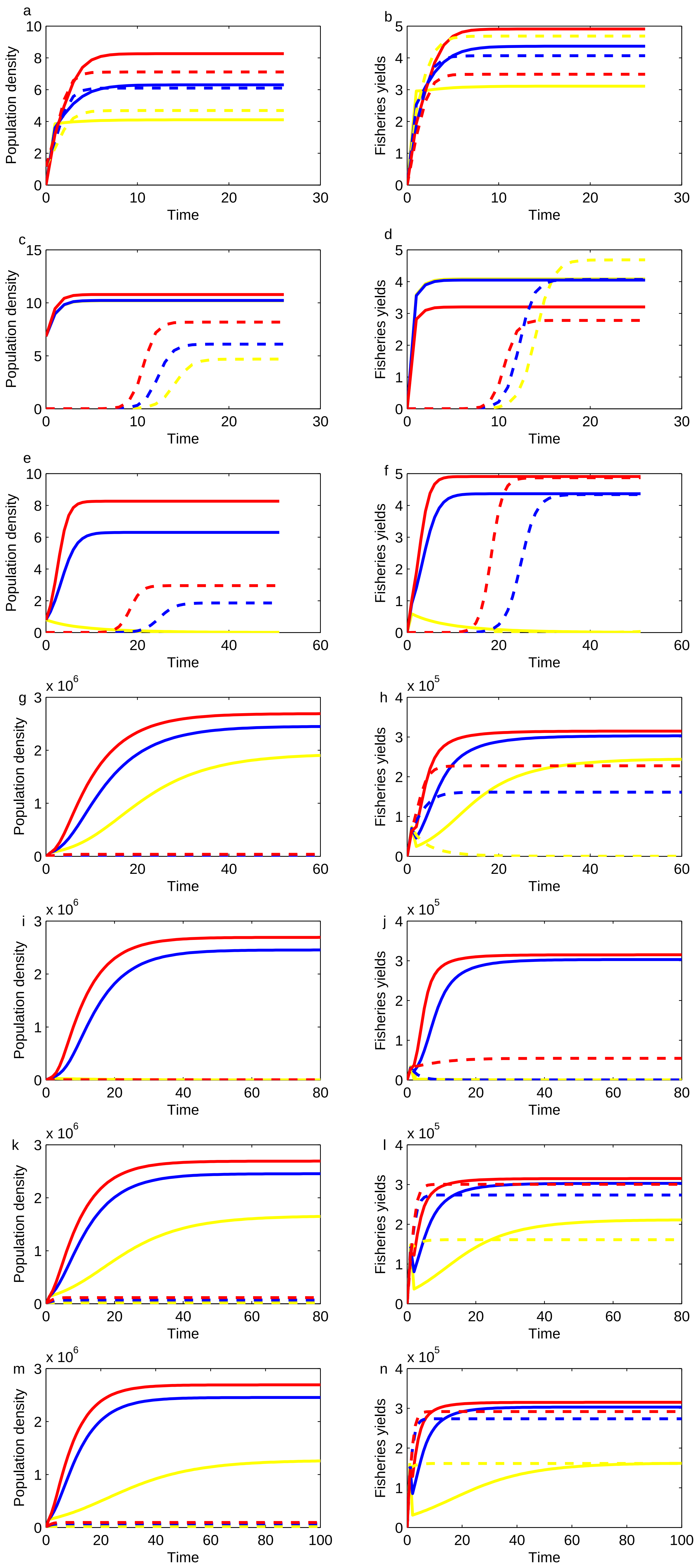
a
b
c
d
e
f
g
h
i
j
k
l
m
n
Population density
Fisheries yields
Time
x 10^6
x 10^5

Fig. S1

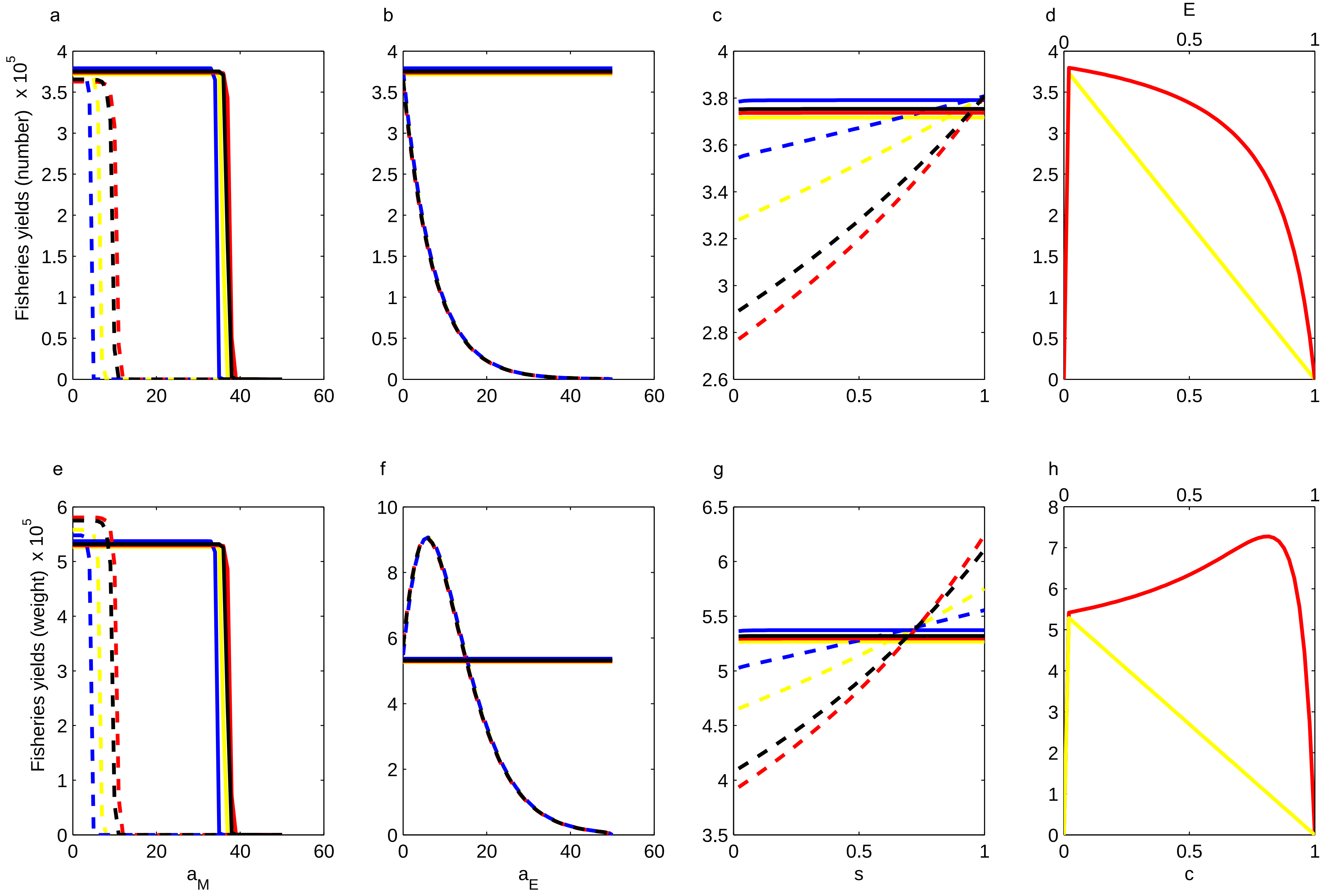
a
b
c
d
e
f
g
h
Fisheries yields (number) x 10^5
Fisheries yields (weight) x 10^5
a_M
a_E
s
c
E

Fig. S2

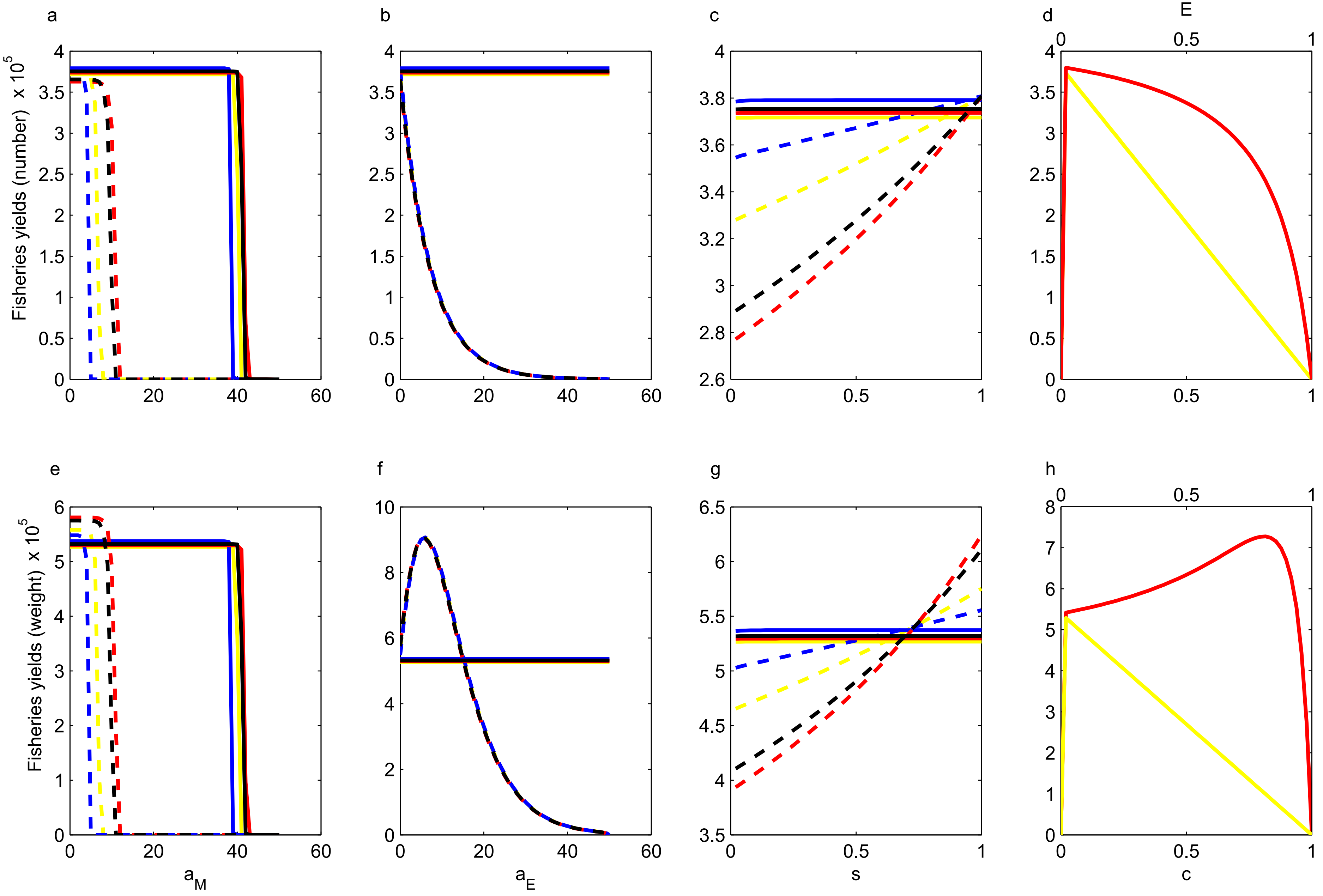
a
b
c
d
E
Fisheries yields (number) x 10^5
e
f
g
h
Fisheries yields (weight) x 10^5
a_M
a_E
s
c

Fig. S3

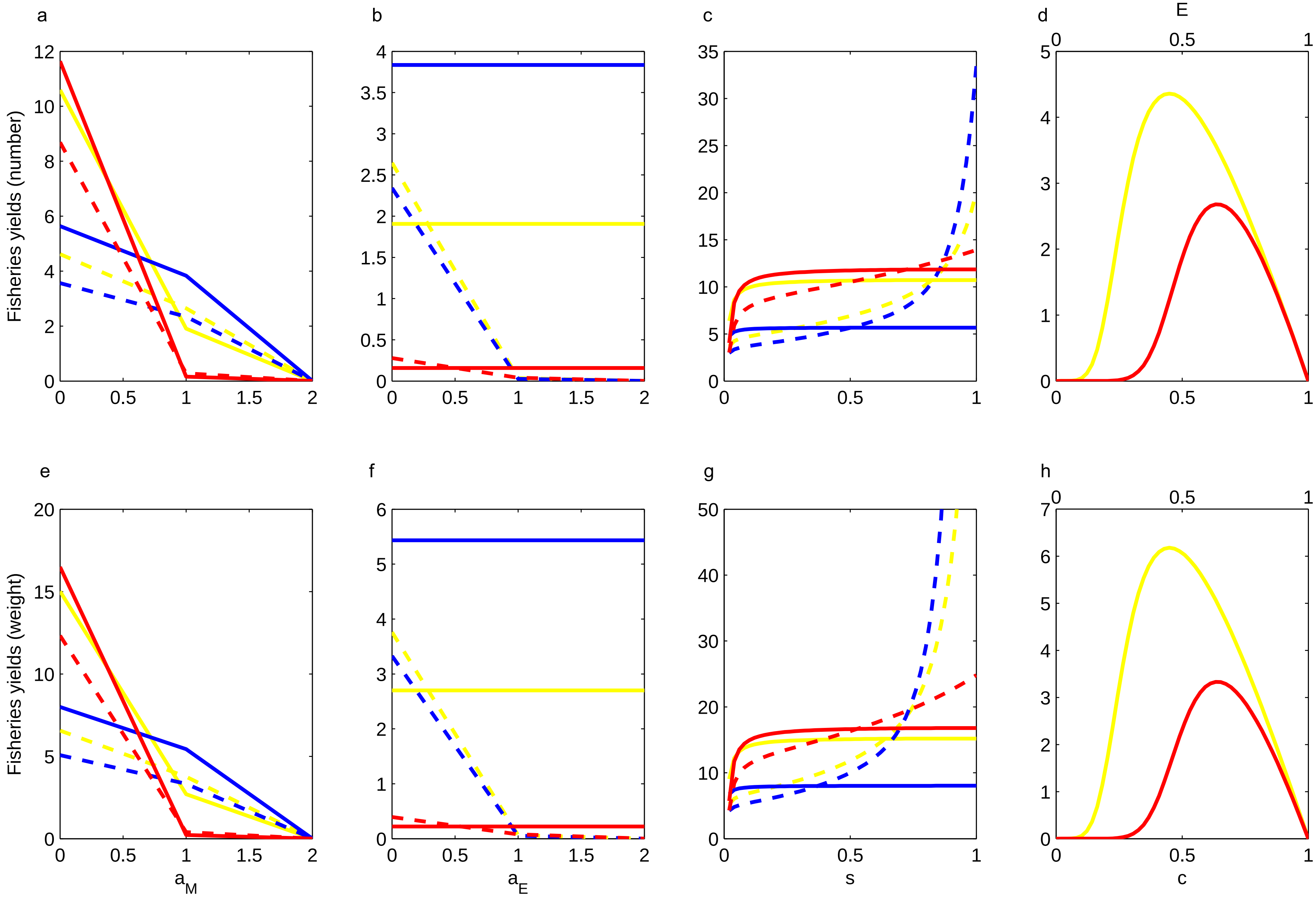
a
b
c
d
E
Fisheries yields (number)
e
f
g
h
Fisheries yields (weight)
$a_M$
$a_E$
s
c

Fig. S4

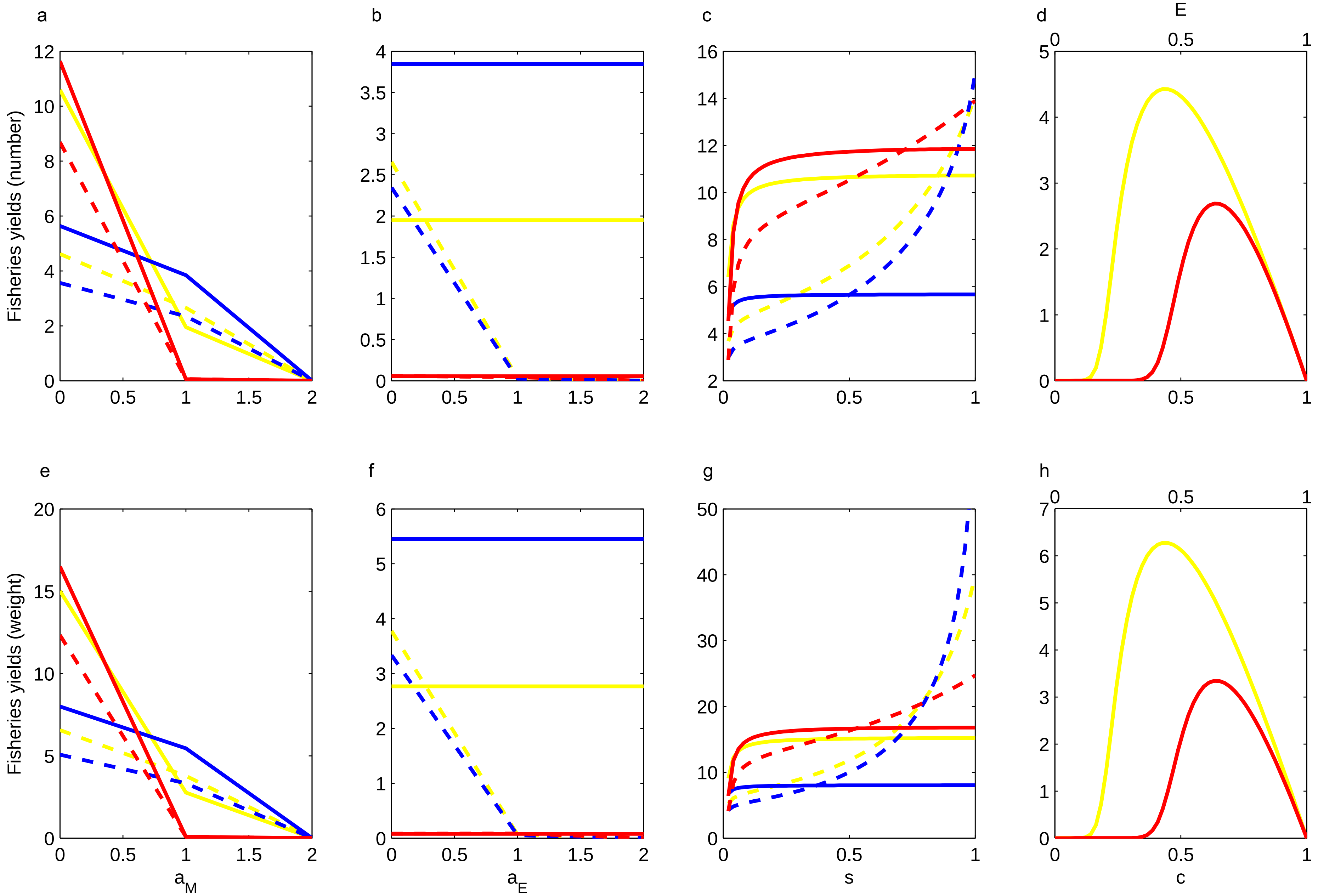
a
b
c
d
E
e
f
g
h
Fisheries yields (number)
Fisheries yields (weight)
$a_M$
$a_E$
s
c

Fig. S5

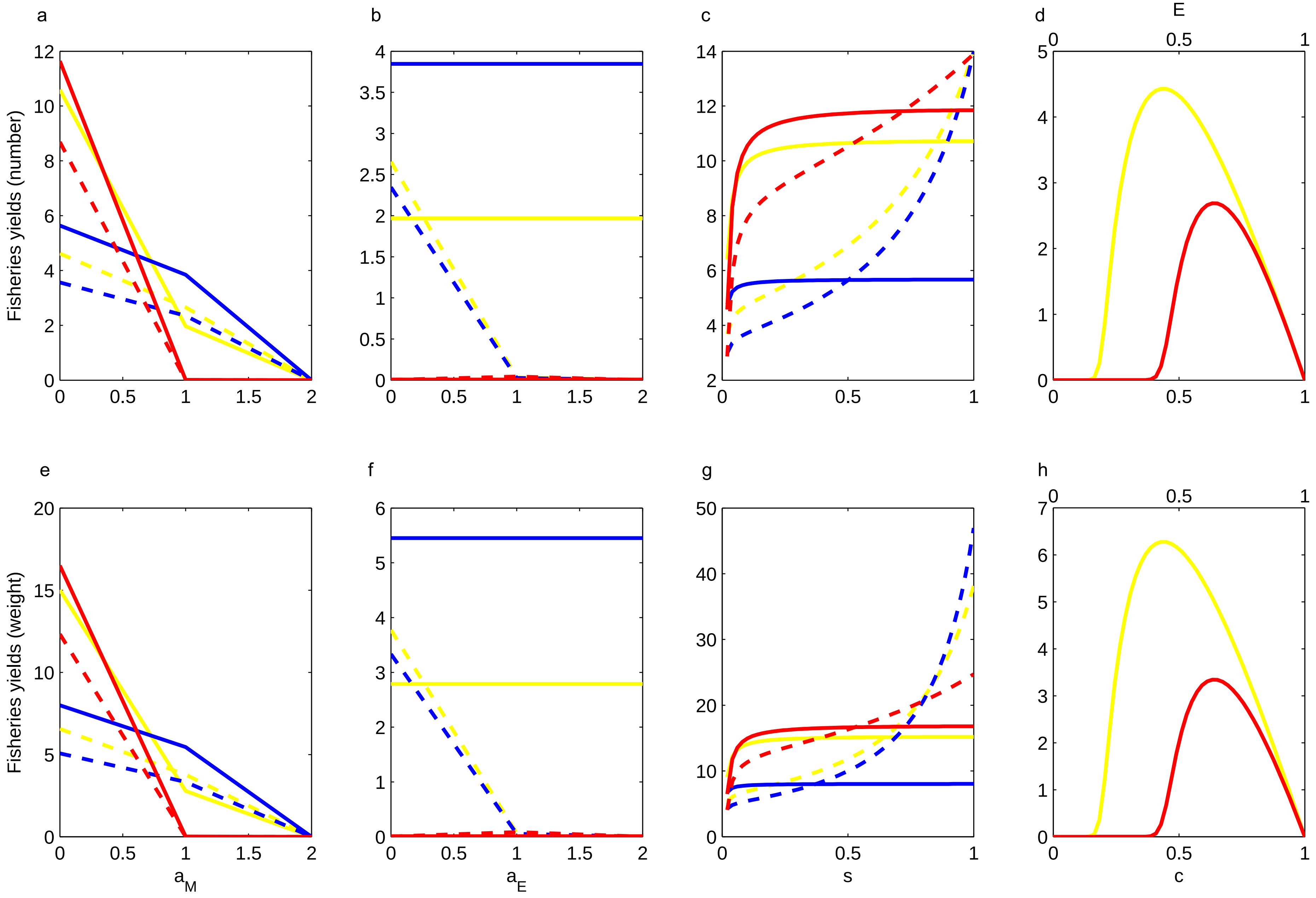

a
b
c
d
e
f
g
h
E
Fisheries yields (number)
Fisheries yields (weight)
$a_M$
$a_E$
s
c


Fig. S6

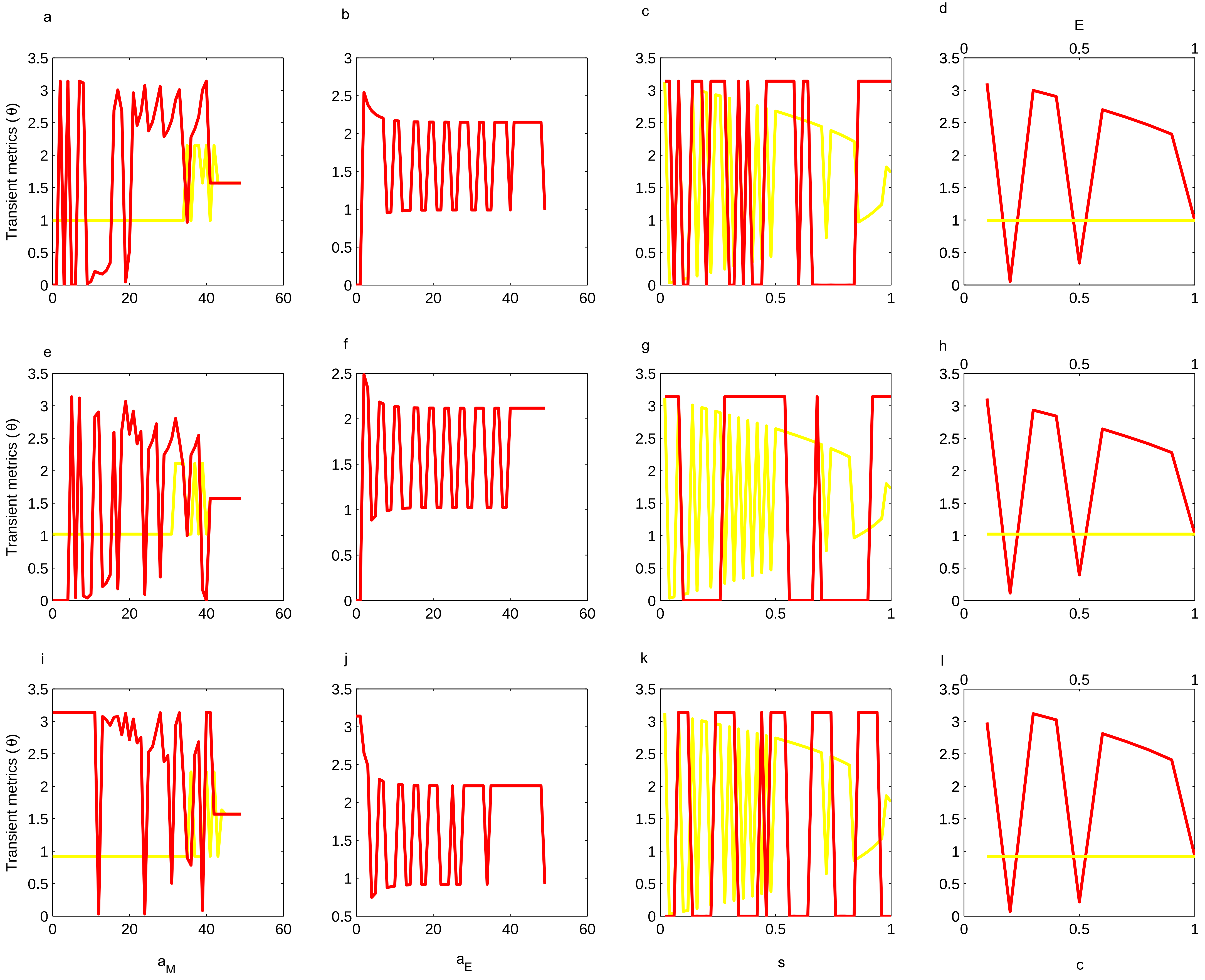
a
b
c
d
E
e
f
g
h
i
j
k
l
Transient metrics (θ)
Transient metrics (θ)
Transient metrics (θ)
a_M
a_E
s
c

Fig. S7

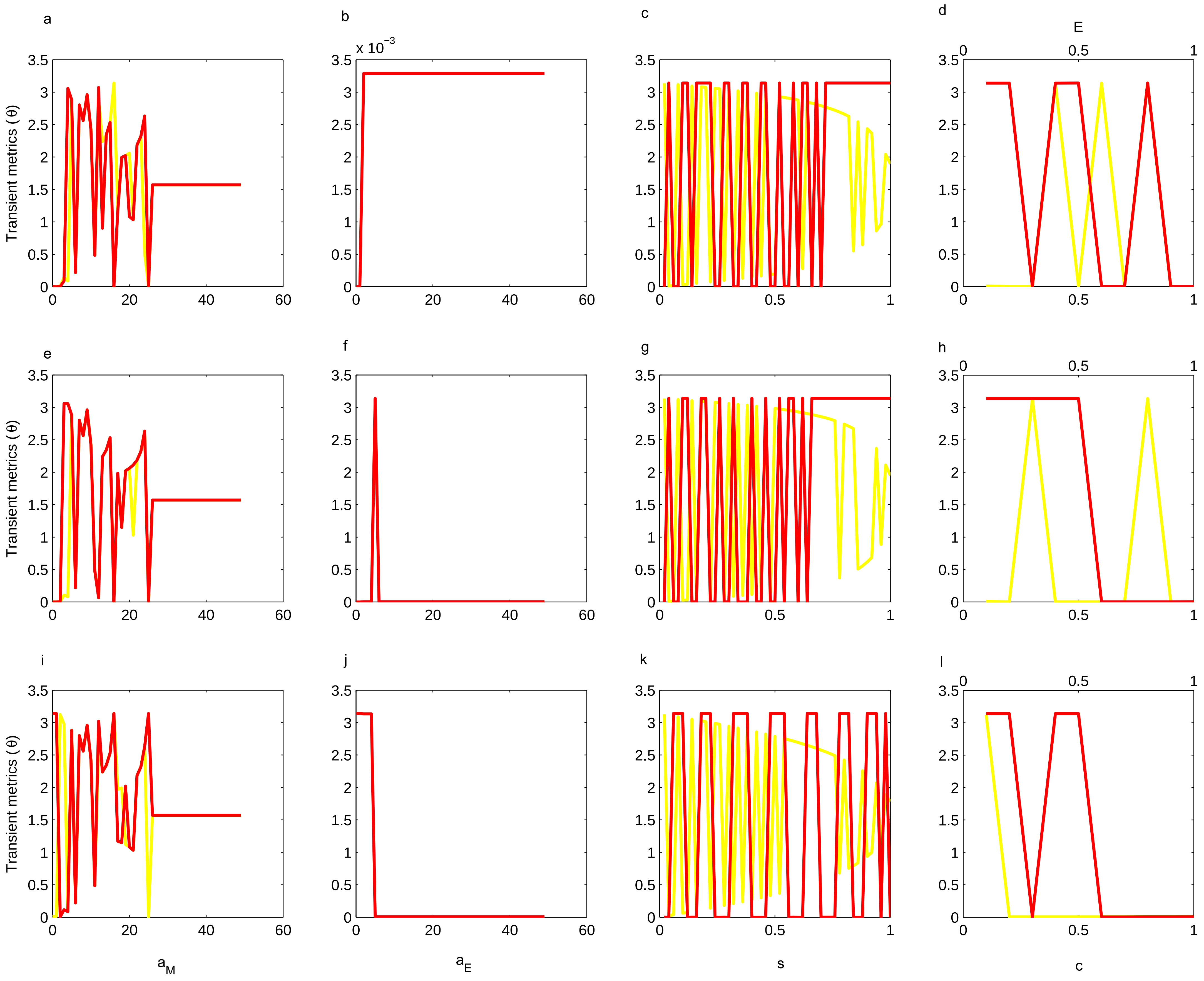


Fig. S8

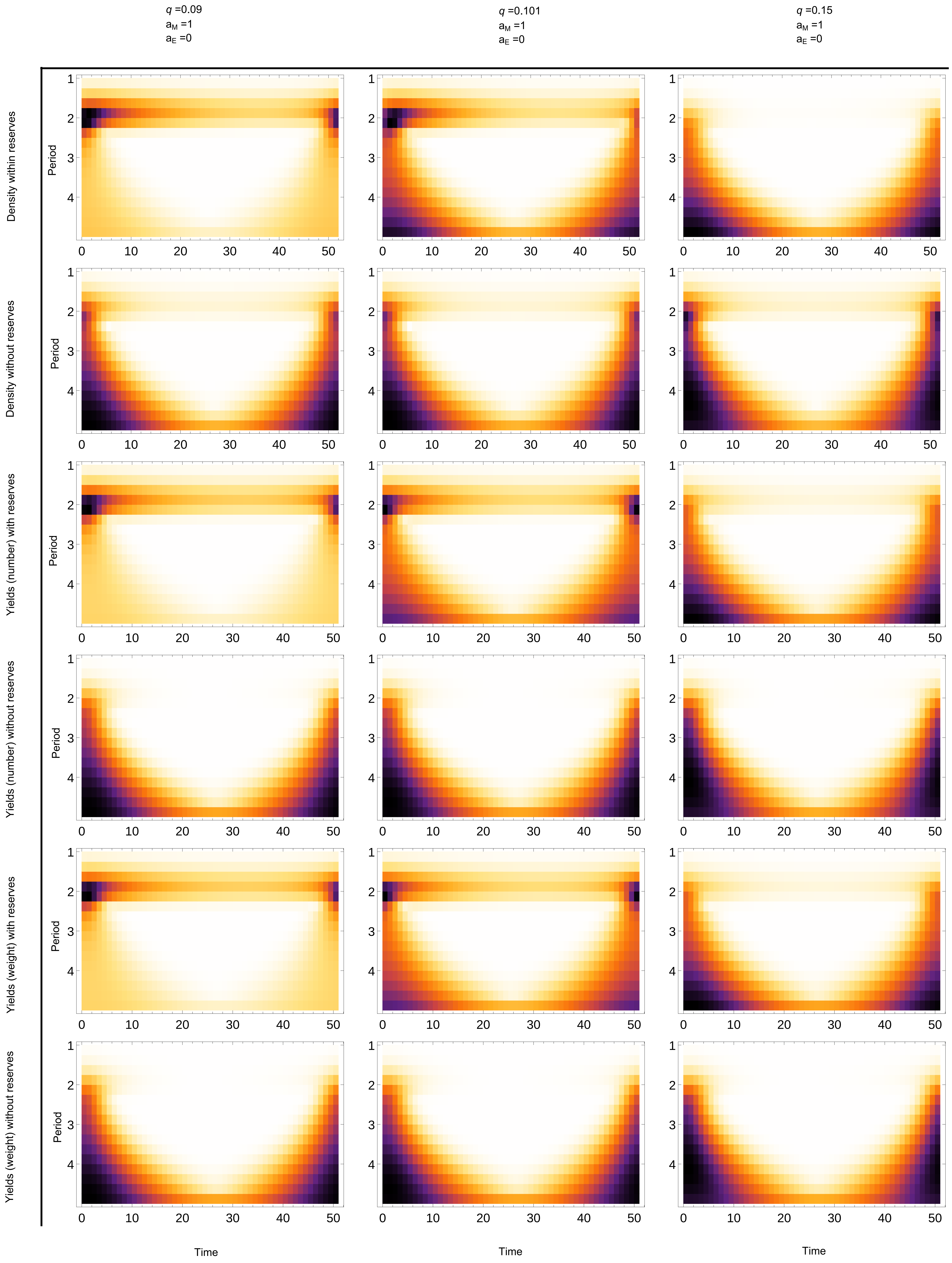

$q$ =0.09
$a_M$ =1
$a_E$ =0
$q$ =0.101
$a_M$ =1
$a_E$ =0
$q$ =0.15
$a_M$ =1
$a_E$ =0
Density within reserves
Density without reserves
Yields (number) with reserves
Yields (number) without reserves
Yields (weight) with reserves
Yields (weight) without reserves
Period
Time


Fig. S9

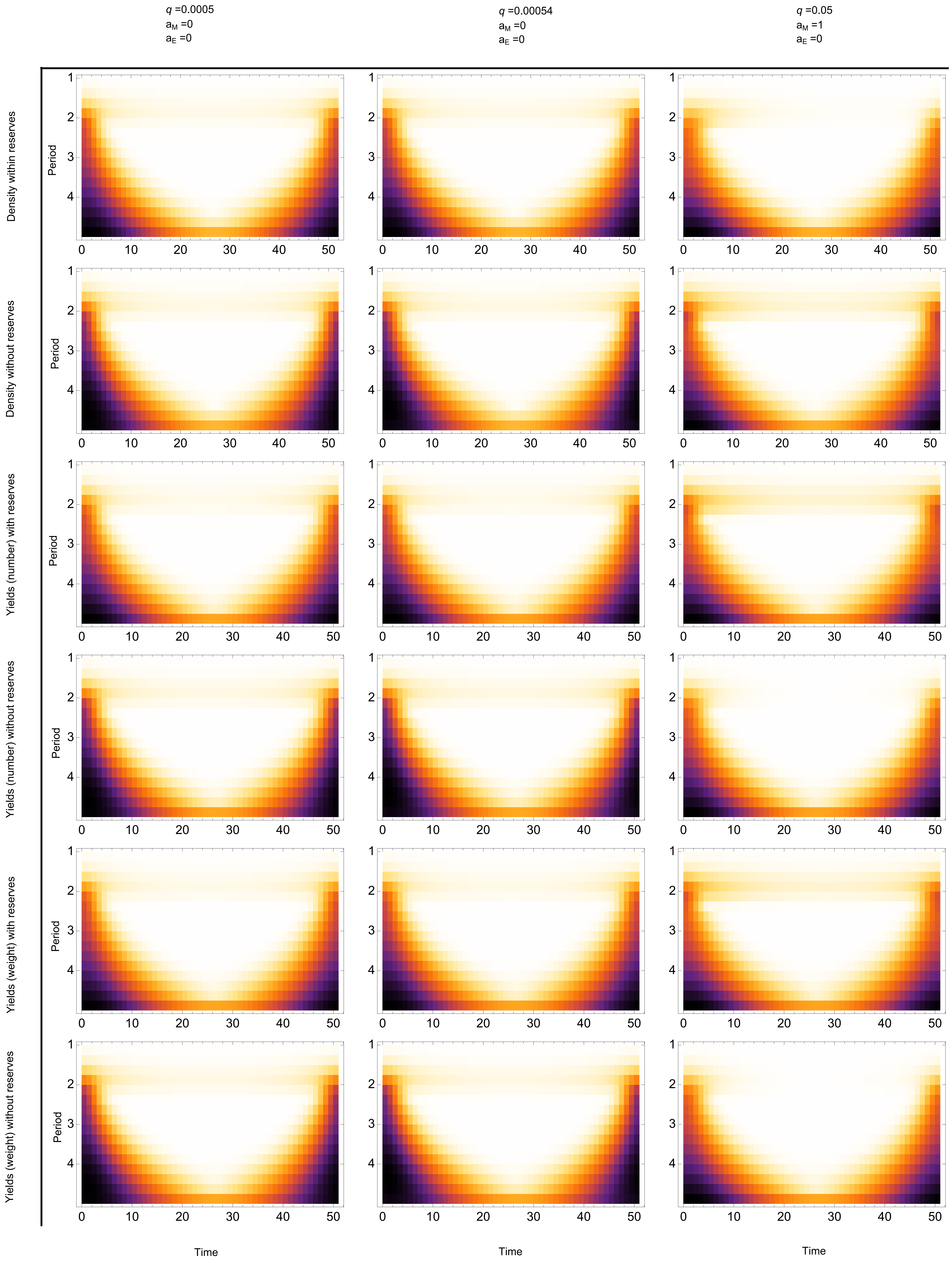

q =0.0005
aM =0
aE =0
q =0.00054
aM =0
aE =0
q =0.05
aM =1
aE =0
Density within reserves
Density without reserves
Yields (number) with reserves
Yields (number) without reserves
Yields (weight) with reserves
Yields (weight) without reserves
Period
Time


Fig. S10

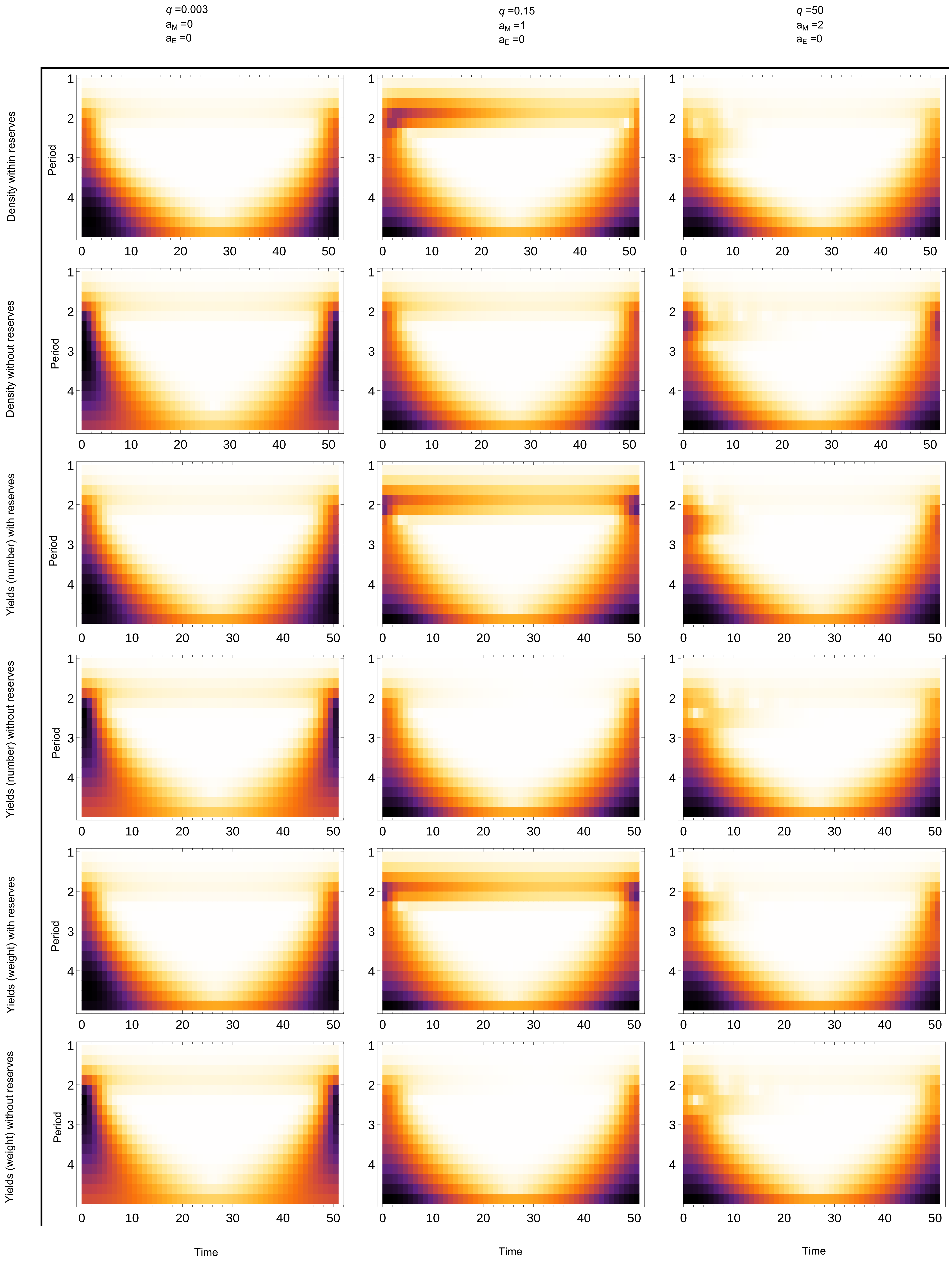

q =0.003
aM =0
aE =0
q =0.15
aM =1
aE =0
q =50
aM =2
aE =0
Density within reserves
Density without reserves
Yields (number) with reserves
Yields (number) without reserves
Yields (weight) with reserves
Yields (weight) without reserves
Period
Time


Fig. S11

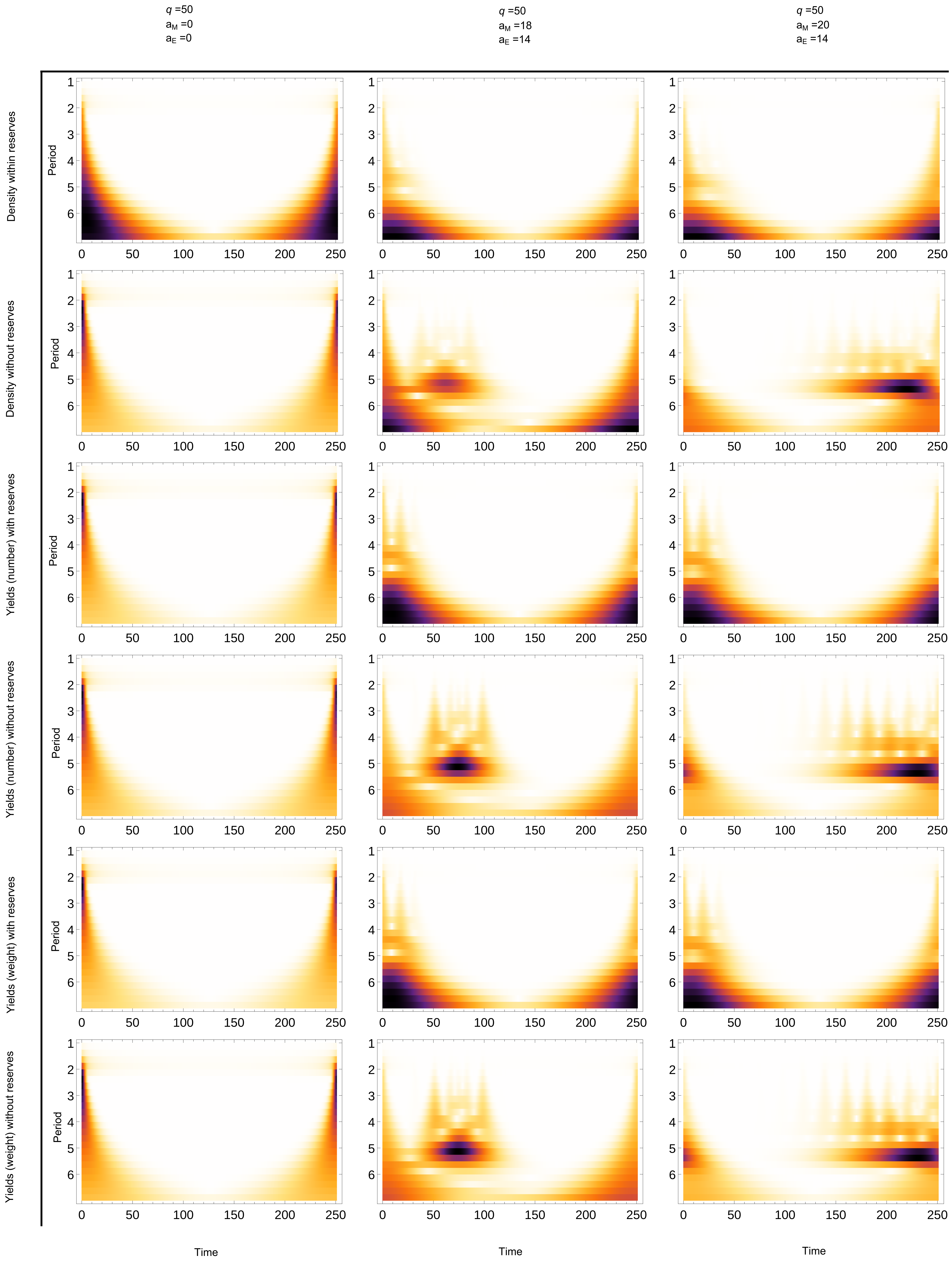

q =50
aM =0
aE =0
q =50
aM =18
aE =14
q =50
aM =20
aE =14
Density within reserves
Density without reserves
Yields (number) with reserves
Yields (number) without reserves
Yields (weight) with reserves
Yields (weight) without reserves
Period
Time


Fig. S12

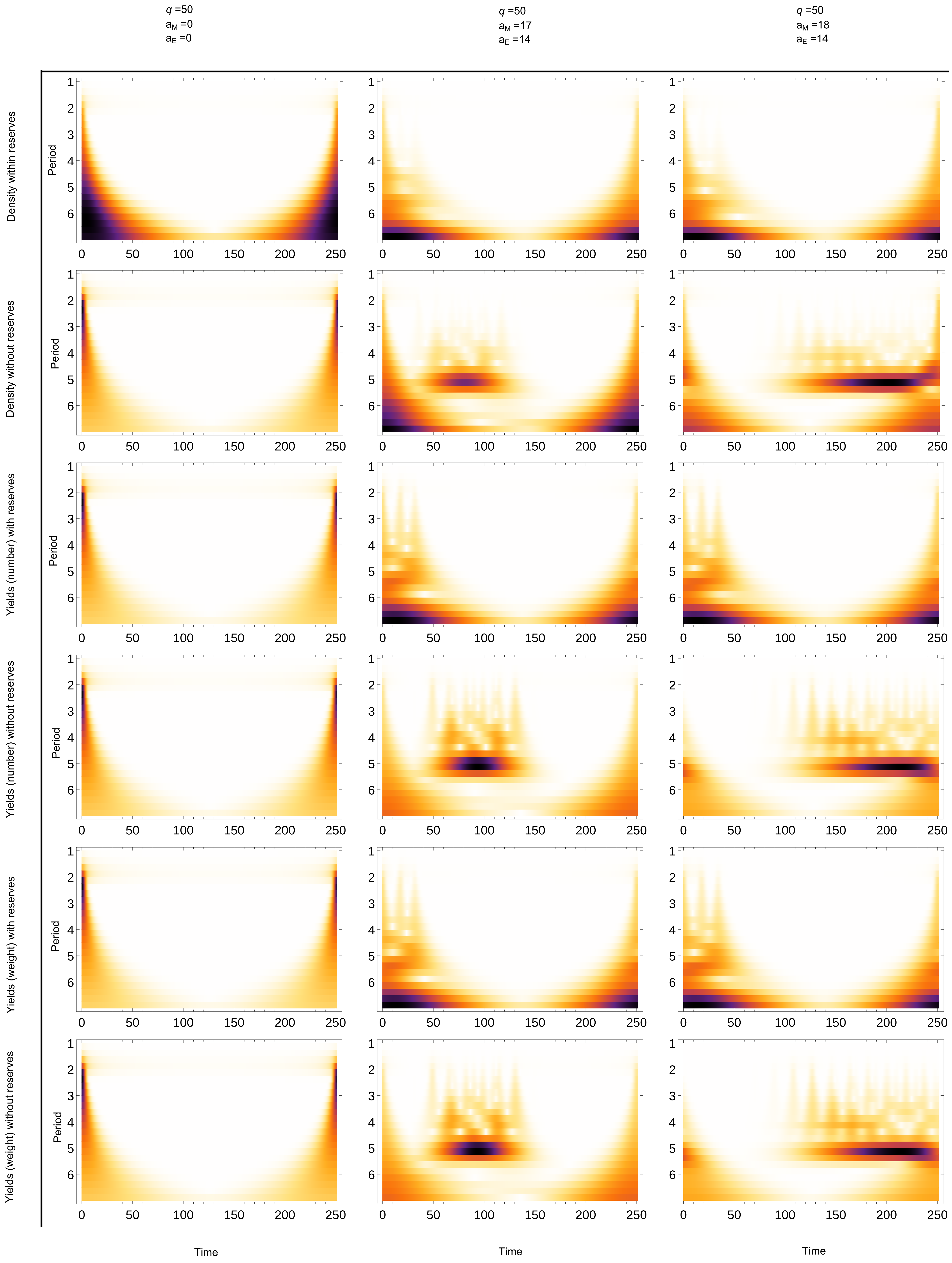

$q$ =50
$a_M$ =0
$a_E$ =0
$q$ =50
$a_M$ =17
$a_E$ =14
$q$ =50
$a_M$ =18
$a_E$ =14
Density within reserves
Density without reserves
Yields (number) with reserves
Yields (number) without reserves
Yields (weight) with reserves
Yields (weight) without reserves
Period
1
2
3
4
5
6
0
50
100
150
200
250
Time


Fig. S13

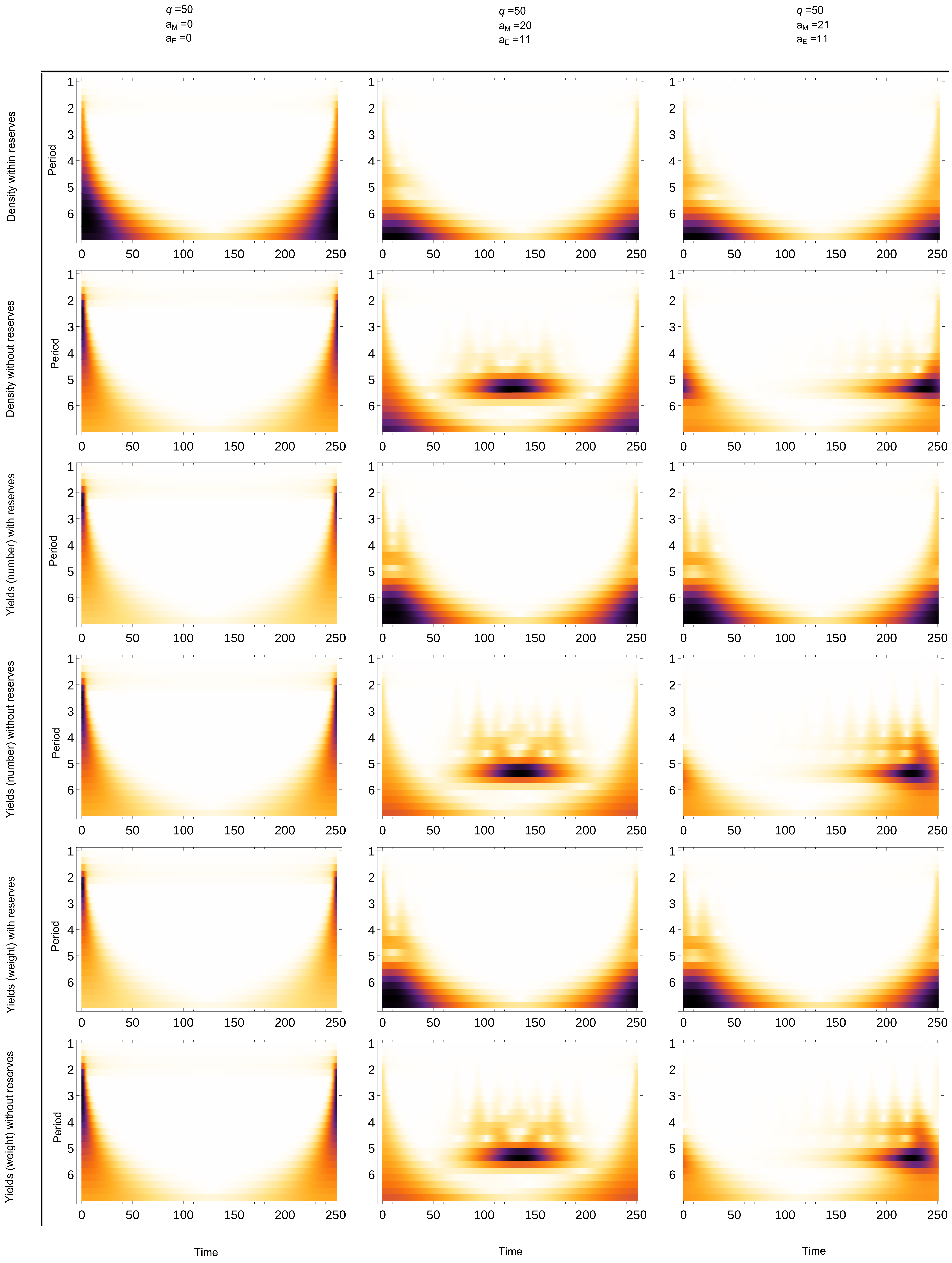

q =50
a_M =0
a_E =0
q =50
a_M =20
a_E =11
q =50
a_M =21
a_E =11
Density within reserves
Density without reserves
Yields (number) with reserves
Yields (number) without reserves
Yields (weight) with reserves
Yields (weight) without reserves
Period
Time


Fig. S14
